# A flexible Bayesian framework for unbiased estimation of timescales

**DOI:** 10.1101/2020.08.11.245944

**Authors:** Roxana Zeraati, Tatiana A. Engel, Anna Levina

## Abstract

Timescales characterize the pace of change for many dynamic processes in nature. Timescales are usually estimated by fitting the exponential decay of data autocorrelation in the time or frequency domain. We show that this standard procedure often fails to recover the correct timescales due to a statistical bias arising from the finite sample size. We develop an alternative approach to estimating timescales by fitting the sample autocorrelation or power spectrum with a generative model based on a mixture of Ornstein-Uhlenbeck processes using adaptive Approximate Bayesian Computations. Our method accounts for finite sample size and noise in data and returns a posterior distribution of timescales that quantifies the estimation uncertainty and can be used for model selection. We demonstrate the accuracy of our method on synthetic data and illustrate its application to recordings from the primate cortex. We provide a customizable Python package implementing our framework with different generative models suitable for diverse applications.

## 1 Introduction

Dynamic changes in many stochastic processes occur over typical periods known as timescales. Timescales of different processes in nature range broadly from milliseconds in protein folding [1, 2] and neural signalling [3–5], to minutes in gene splicing [6–8] and synaptic plasticity [9–12], to days in spreading of infectious diseases [13–15], to years in demographic changes of populations [16, 17], and up to millennia in the climate change [18, 19]. Accurate measurements of timescales from experimental data are necessary to uncover mechanisms controlling the dynamics of underlying processes and reveal their function. The variation of timescales across brain areas reflects the functional hierarchy in the neocortex and areal differences in temporal integration of information [3, 20–22], while aberrant timescales were associated with autism [23]. Precise estimation methods are necessary to detect changes in timescales during cognitive processes such as attention [24]. Estimated timescales are also used to determine the operating regime of the dynamics, e.g., how close the system is to a critical point [25], which can reveal general principles organizing the collective behavior of complex systems such as the brain [26–28]. Thus, many problems require estimating timescales from experimental data to constrain theoretical models and enable accurate predictions in practical applications.

The timescales of a stochastic process are defined by the exponential decay rates of its autocorrelation function. Accordingly, timescales are usually estimated by fitting the autocorrelation of a sample time-series with exponential decay functions [3,25,29–40] or using the autocorrelation half-width [20, 23, 41]. Equivalently, timescales can be estimated in frequency domain by fitting the shape of the sample power spectral density (PSD) with a Lorentzian function which is the PSD of a process with an exponentially decaying autocorrelation [22,42]. However, many factors unrelated to the dynamics of the processes under study can alter the shape of autocorrelations and, according to the Wiener–Khinchin theorem [43], the PSD shape. For example, autocorrelations of *in vivo* neural activity can contain components arising from a specific trial structure of a behavioral task or slow drifts in the average activity. To correct for these irrelevant factors, several techniques were developed based on data resampling, in particular, trial shuffling and spike jittering methods [44–47]. These methods remove from the autocorrelation the average autocorrelation of surrogate data, which are designed to match the irrelevant factors in the real data but are otherwise random. The success of these methods critically depends on the ability to construct the appropriate surrogate data that exactly reproduce the irrelevant factors.

The values of sample autocorrelation computed from a finite time series systematically deviate from the true autocorrelation [48–53]. This bias is exacerbated by the lack of independence between the samples, which is particularly harmful when the timescales are large and timeseries are short [54, 55]. Generally, the magnitude of bias depends on the length of the sample time-series, but also on the value of the true autocorrelation at each time-lag. The expected value and variance of the autocorrelation bias can be derived analytically in some simple cases, such as a Markov process with a single timescale [48, 51]. However, the analytical derivation is not tractable for more general processes that involve multiple timescales or have additional temporal structure. Moreover, since the bias depends on the true autocorrelation itself, which is unknown, it cannot be corrected by constructing appropriate surrogate data as in shuffling or jittering methods.

The statistical bias deforms the shape of empirical autocorrelations and PSDs and hence can affect the timescales estimated by direct fitting of the shapes. Indeed, it was noticed that fitting the sample autocorrelation of an Ornstein-Uhlenbeck (OU) process with an exponential decay function results in systematic errors in the estimated timescale and the confidence interval [56]. To avoid these errors, it is possible to fit the time-series directly with an autoregressive model, without using autocorrelation or PSD [21, 56]. For example, the parameters of an OU process can be obtained with a maximum-likelihood estimator, which directly fits time-series and thus evades the autocorrelation bias [56]. However, the advantage of autocorrelation is that irrelevant factors (e.g., slow activity drifts) can be efficiently removed with resampling methods (e.g., spike jittering), and any additional dynamics not locked to trial onset (e.g., oscillations) can be easily incorporated in the autocorrelation shape. In contrast, accounting for irrelevant factors and dynamics in the raw time-series is generally difficult as it requires adding components to the autoregressive model matching the temporal structure of these factors on single trials, all of which need to be fitted to the data. Thus, fitting the summary statistic such as autocorrelation or PSD is attractive, but how the statistical bias affects the estimated timescales was not studied systematically. Moreover, rather than maximum-likelihood estimates, inference of a full posterior distribution is often necessary, for example, for model selection, which has not been addressed for complex processes with multiple timescales, additional temporal structure and noise.

We show that large systematic errors in estimated timescales arise from the statistical bias due to a finite sample size, which is evident in different processes with various number of timescales and different trial durations. These estimation errors may result in misleading interpretations of the underlying biological processes. To correct for the bias, we develop a flexible computational framework based on adaptive Approximate Bayesian Computations (aABC) that estimates timescales by fitting the autocorrelation or PSD with a generative model. ABC is a family of likelihood-free inference algorithms for estimating model parameters when the likelihood function cannot be calculated analytically [57]. The aABC algorithm approximates the multivariate posterior distribution of parameters of a generative model using population Monte-Carlo sampling [58]. Our generative model is based on a mixture of Ornstein-Uhlenbeck processes—one for each estimated timescale—which have exponentially decaying autocorrelations. The generative model can be further augmented with the desired noise model (e.g., a spike generation process) and additional temporal structure to match the statistics of the data. Our method accurately recovers the correct timescales from finite data samples for various synthetic processes with known ground-truth dynamics. The inferred posterior distributions quantify the uncertainty of estimates and can be used for model selection to compare alternative hypotheses about the dynamics of the underlying process. To illustrate an application of our method, we estimate timescales of ongoing spiking activity in the primate visual cortex during a behavioral task. The computational framework presented here can be adapted to various types of data and can find broad applications in neuroscience, cellular biology, epidemiology, physics, and other fields. To allow for an easy adoption and further development of our framework, we provide a Python package called abcTau that supports different types of data and generative models, and includes the model comparison functionality and a parallel processing option for analyzing large-scale datasets.

## 2 Results

### 2.1 Bias in timescales estimated by direct fitting

Timescales of a stochastic process *A*(*t′*) are defined by the exponential decay rates of its autocorrelation function. The autocorrelation is the correlation between the values of the process at two time points separated by a time lag *t*. For stationary processes, the autocorrelation function only depends on the time lag:

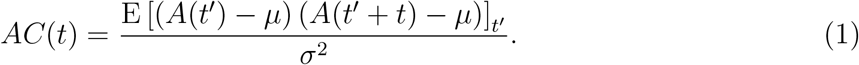

Here *μ* and *σ*^2^ are, respectively, the mean and variance of the process, which are constant in time, and E[·]_*t′*_ is the expectation over *t′*. Different normalizations of autocorrelation are used in literature, but our results do not depend on a specific choice of normalization.

In experiments or simulations, the autocorrelation needs to be estimated from a finite sample of empirical data. A data sample from the process *A*(*t′*) constitutes a finite time-series measured at discrete times 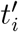 (*i* = 1… *N*, where *N* is the length of the time-series). For example, the sample time-series can be spike-counts of a neuron in discrete time bins, or a continuous voltage signal measured at a specific sampling rate. Accordingly, the sample autocorrelation is defined for a discrete set of time lags *t_j_*. For empirical data, the true values of *μ* and *σ* are unknown. Hence, several estimators of the sample autocorrelation were proposed, which use different estimators for the sample mean 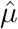 and sample variance 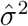 [48, 49]. One possible choice is:

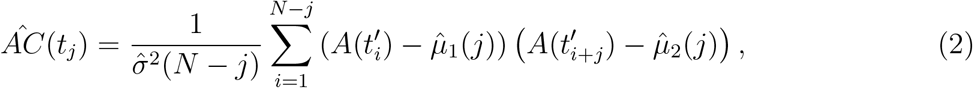

with the sample variance 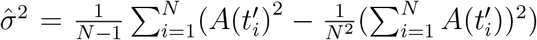 and two different sample means 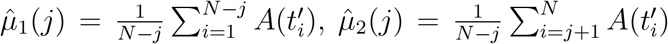. The sample autocorrelation can also be computed as the inverse Fourier transform of PSD, based on the Wiener–Khinchin theorem [43]. However, for any of these methods, the sample autocorrelation is a biased estimator: for a finite length time-series the values of the sample autocorrelation systematically deviate from the ground-truth autocorrelation [48–53] (Fig. 1, Supplementary Fig. 1). This statistical bias deforms the shape of the sample autocorrelation or PSD and therefore may affect the estimation of timescales by direct fitting of the shape.

**Fig. 1.**
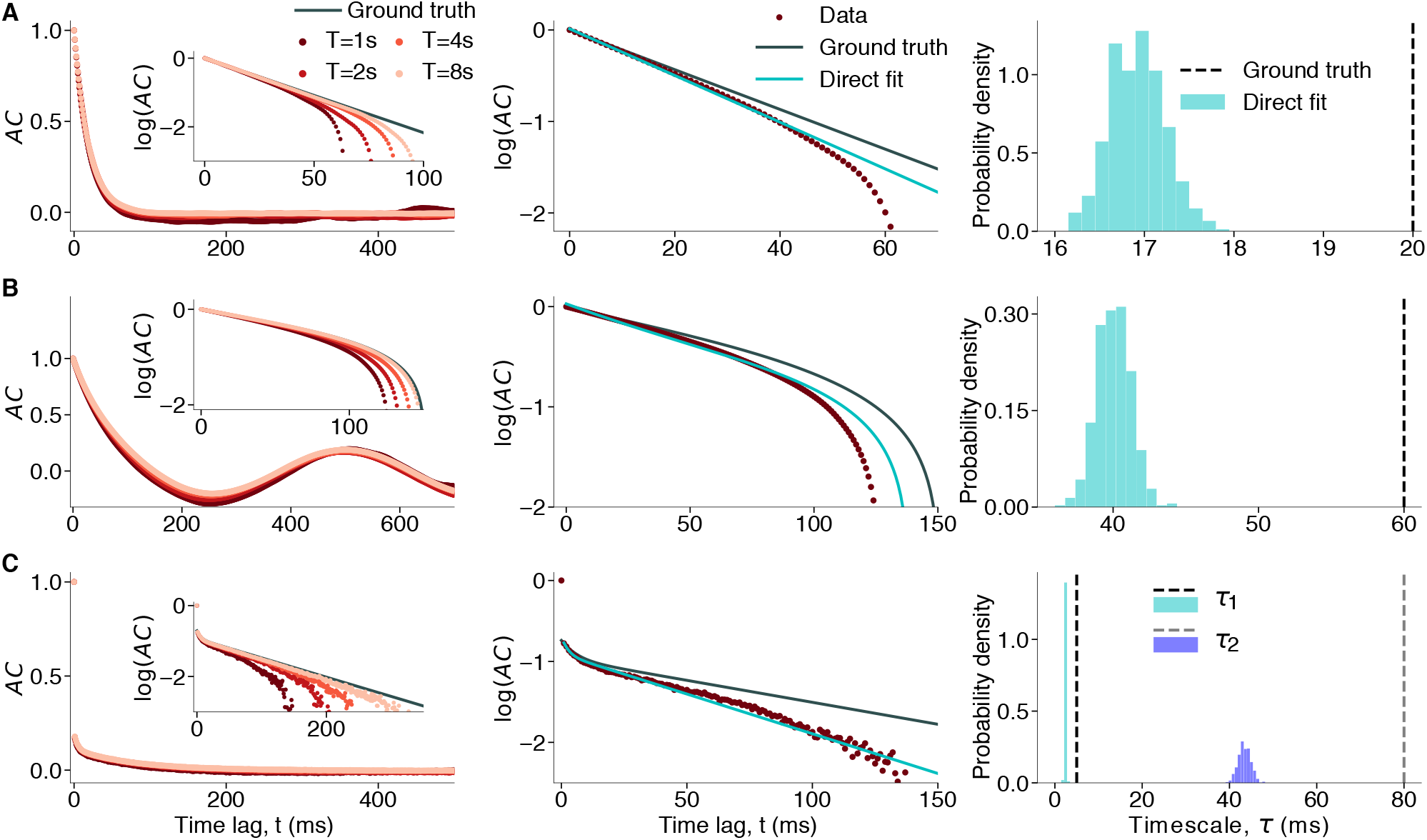
Bias in sample autocorrelations and timescales estimated by direct exponential fitting. **(a)** Data are generated from an OU process with the timescale *τ* = 20 ms. **Left**: Sample autocorrelation (colored dots) systematically deviates from the ground-truth autocorrelation (gray line). The shape of sample autocorrelation depends on the time-series duration (*T*) approaching the ground truth with increasing *T* (inset: close up in logarithmic-linear scale). **Middle:** Direct fitting of the analytical autocorrelation function (cyan) to sample autocorrelation (brown, *T* = 1 s) does not recover the ground-truth autocorrelation shape (gray). **Right:** The ground-truth timescales largely deviate from the distribution of timescales estimated by direct exponential fitting across 500 independent realizations of the same process. **(b)** Same as a for data from a linear mixture of an OU process with the timescale *τ* = 60 ms and an oscillation with frequency *f* = 2 Hz (Eq. 5). **(c)** Same as a for data from an inhomogeneous Poisson process with the instantaneous rate generated from a linear mixture of two OU processes with timescales *τ*_1_ = 5 ms and *τ*_2_ = 80 ms. All simulation parameters are provided in Methods 4.5.

To investigate how the autocorrelation bias affects the timescales estimated by direct exponential fitting, we tested how accurately this procedure recovers the correct timescales on synthetic data with a known ground truth. We generated synthetic data from several stochastic processes for which the autocorrelation function can be computed analytically. The exponential decay rates of the analytical autocorrelation provide the ground-truth timescales. Each synthetic dataset consisted of 500 independent realizations of the process (i.e. trials) with fixed duration. Such trial-based data are typical in neuroscience but usually with a smaller number of trials. We computed the sample autocorrelation for each trial using Eq. 2, averaged them to reduce the noise, and then fitted the average autocorrelation with the correct analytical functional form to estimate the timescale parameters. We repeated the entire procedure 500 times to obtain a distribution with multiple independent samples of timescales estimated by direct fit (i.e. we simulated 500 × 500 trials for each distribution).

We considered three ground-truth processes which differed in the number of timescales, additional temporal structure and noise. First, we used an Ornstein–Uhlenbeck (OU) process (Fig. 1A):

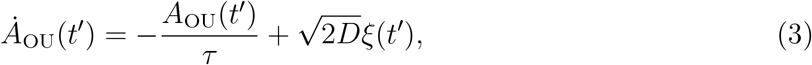

where *ξ*(*t′*) is a Gaussian white noise with zero mean, and the diffusion parameter *D* sets the variance Var[A_OU_(*t′*)] = *D_τ_* [59, 60]. The autocorrelation of the OU process is an exponential decay function [61]

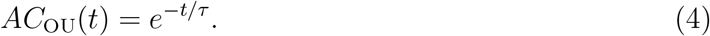

Accordingly, the parameter *τ* provides the ground-truth timescale.

Second, we used a linear mixture of an OU process and an additional oscillatory component (Fig. 1B):

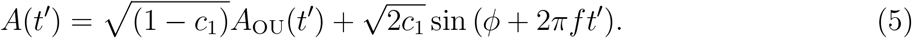

Here the coefficient *c*_1_ ∈ [0, 1] sets the relative weights of two components without changing the total variance of the process, and the phase *ϕ* is drawn on each trial from a uniform distribution on [0, 2*π*]. This process resembles the firing rate of a neuron modulated by a slow oscillation [62, 63]. The autocorrelation of this process is given by

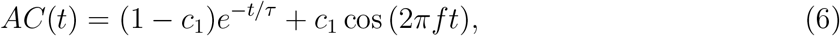

hence, the ground-truth timescale is defined by the OU parameter *τ*.

Finally, we used an inhomogeneous Poisson process with the instantaneous rate modeled as a linear mixture of two OU processes with different timescales (Eq. 13, Fig. 1C). The OU mixture was shifted, scaled and rectified to produce the Poisson rate with the desired mean and variance (Eq. 17). Similar doubly-stochastic processes are often used to model spiking activity of neurons [64]. The total variance of this process consists of two contributions: the rate variance and the Poisson process variance (Eq. 19) [65]. We simulated this process in discrete time bins 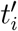 by sampling event-counts from a Poisson distribution with the given instantaneous rate (Methods 4.1.3). The autocorrelation at all lags *t_j_* (*j* > 0) arises only from the autocorrelation of the rate, since the Poisson process has no temporal correlations. The drop of autocorrelation between *t*_0_ and *t*_1_ mainly reflects the Poisson-process variance, and for all non-zero lags *t_j_* (*j* > 0) the autocorrelation is given by

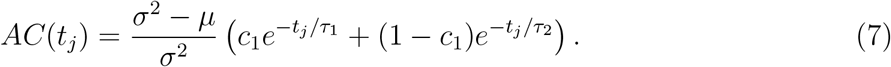

Here *τ*_1_ and *τ*_2_ are the ground-truth timescales defined by the parameters of two OU processes, *c*_1_ is the mixing coefficient, and *μ* and *σ*^2^ are the mean and variance of the event-counts, respectively.

For all three processes, the sample autocorrelation exhibits a negative bias: the values of the sample autocorrelation are systematically below the ground-truth autocorrelation function (Fig. 1, left). This bias is clearly visible in the logarithmic-linear scale, where the ground-truth exponential decay turns into a straight line. The sample autocorrelation deviates from the straight line and even becomes systematically negative at intermediate lags (hence not visible on the logarithmic scale) for processes with a strictly positive ground-truth autocorrelation (Fig. 1a,c). The deviations from the ground truth are larger when the timescales are longer or when multiple timescales are involved. The negative bias decreases for longer trial durations (Fig. 1, left inset), but it is still substantial for realistic trial durations such as in neuroscience data.

Due to the negative bias, a direct fit of the sample autocorrelation with the correct analytical function cannot recover the ground-truth timescales (Fig. 1, middle, right, Supplementary Fig. 2, 3). When increasing the duration of each trial, the timescales obtained from the direct fits become closer to the ground-truth values (Supplementary Fig. 4). This observation indicates that timescales estimated from datasets with different trial durations cannot be directly compared, as differences in the estimation bias may result in misleading interpretations. Thus, direct fitting of a sample autocorrelation, even when the correct analytical form of autocorrelation is known, is not a reliable method for measuring timescales in experimental data. Bias-correction methods based on parametric bootstrapping can mitigate the estimation bias of timescales from direct fits [66]. However, since the amount of bias depends on the ground-truth timescales, such corrections cannot guarantee accurate estimates in all cases (Supplementary Fig. 5).

Alternatively, we can estimate the timescales by fitting the PSD shape in the frequency domain. In this approach, the PSD is fitted with a Lorentzian function [22,42] which is the ground-truth PSD for a stochastic process with an exponentially decaying autocorrelation (e.g., OU process)

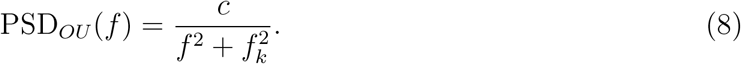

Here *f* is the frequency and 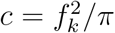 is the normalization constant. From the knee frequency *f_k_*, we can estimate the timescale as *τ* = (2*πf_k_*)^-1^. Comparison of the ground-truth PSD with the sample PSD of an OU process with a finite duration reveals that a statistical bias observed in the time domain also persists in the frequency domain (Supplementary Fig. 6A). Due to this bias, the estimated knee frequency deviates from the ground-truth knee frequency depending on the fitted frequency range, which results in biased estimation of the timescale. Although careful choice of the fitted frequency range can improve the accuracy of estimates (Supplementary Fig. 6B), without knowing the ground-truth timescale, there is no principled way to choose the correct frequency range that will work in all cases. Slightly changing the fitted frequency range can produce large errors in estimated timescale, especially in the presence of additional noise e.g., in spiking activity.

### 2.2 Estimating timescales by fitting generative models with Approximate Bayesian Computations

Since direct fitting of the sample autocorrelation (or PSD) cannot estimate timescales reliably, we developed an alternative computational framework based on fitting the sample autocorrelation (or PSD) with a generative model. Using a generative model with known ground-truth timescales, we can generate synthetic data matching the essential statistics of the observed data, i.e. with the same duration and number of trials, mean and variance. Hence the sample autocorrelations (or PSDs) of the synthetic and observed data will be affected by a similar statistical bias when their shapes match. As a generative model, we chose a linear mixture of OU processes—one for each estimated timescale—which if necessary can be augmented with additional temporal structure (e.g., oscillations) and noise. The advantage of using a mixture of OU processes for modelling the observed data is that the analytical autocorrelation function of this mixture explicitly defines the timescales. We set the number of components in the generative model in accordance with our hypothesis about the shape of the autocorrelation in the data, e.g., the number of timescales, additional temporal structure and noise (Methods 4.1). Then, we optimize parameters of the generative model to match the shape of the autocorrelation (or PSD) between the synthetic and observed data. The timescales of the optimized generative model provide an approximation for the timescales in the observed data, without the bias corruption.

For complex generative models, calculating the likelihood can be computationally expensive or even intractable. Therefore, we optimize the generative model parameters using adaptive Approximate Bayesian Computations (aABC) [58] (Fig. 2). aABC is an iterative algorithm that minimizes the distance between the summary statistic of synthetic and observed data. Depending on the application, we can choose a different summary statistic in time or frequency domain. For example, we can use autocorrelations as the summary statistic and define the suitable distance *d* between the autocorrelations of synthetic and observed data, e.g., as

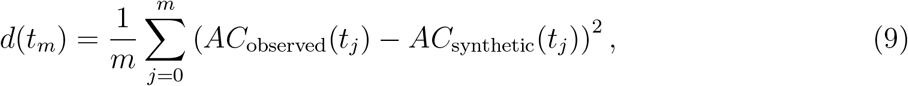

where *t_m_* is the maximum time-lag considered in computing the distance. Alternatively, we can compute distances between the PSDs of synthetic and observed data:

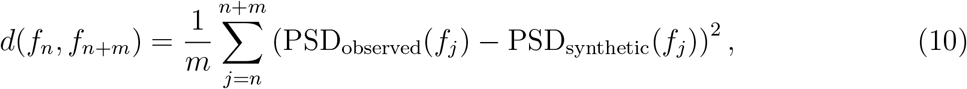

where *f_n_* and *f_n+m_* define the frequency range for computing the distance.

Due to stochasticity of the observed data, point estimates (i.e. assigning a single value to a timescale) are not reliable as different realizations of the same stochastic process lead to slightly different autocorrelation shapes (Supplementary Fig. 7). aABC overcomes this problem by estimating the joint posterior distribution of the generative model parameters, which quantifies the estimation uncertainty.

**Fig. 2.**
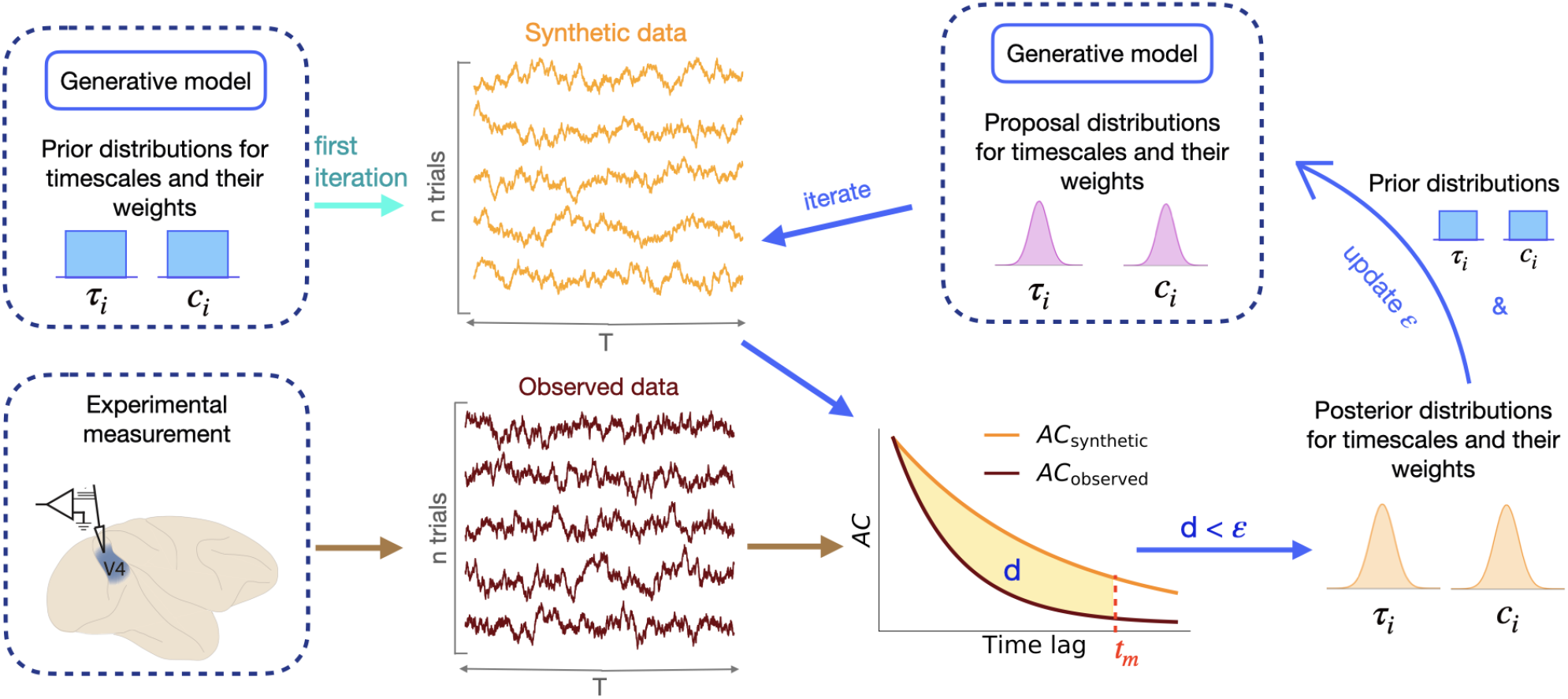
Estimation of timescales with adaptive Approximate Bayesian Computations. aABC estimates timescales by fitting the sample autocorrelation of observed data with a generative model. At the first iteration of the algorithm, parameters of the generative model are drawn from the multivariate prior distribution, e.g., a uniform distribution (upper left). Synthetic data are generated from the generative model with these parameters. If the distance *d* between the autocorrelations of synthetic and observed data is smaller than a specific error threshold *ε*, these parameters are added to the multivariate posterior distribution (lower right). In the subsequent iterations, new parameters are drawn from a proposal distribution which is computed based on the posterior distribution of the previous iteration and the initial prior distribution (upper right, see Methods 4.2 and 4.3).

The aABC algorithm operates iteratively (Fig. 2). First, we choose a multivariate prior distribution over the parameters of the generative model and set an initial error threshold *ε* at a rather large value. On each iteration, we draw samples of the parameter-vector *θ*. On the first iteration, *θ* is drawn directly from the prior distribution. On subsequent iterations, *θ* is drawn from a proposal distribution defined based on the prior distribution and the parameter samples accepted on the previous iteration. We use the generative model with the sample parameters *θ* to generate synthetic data. If the distance *d* between the summary statistic (autocorrelation or PSD) of the observed and synthetic data (Eq. 9, 10) is smaller than the error threshold, the sample *θ* is accepted. We then repeatedly draw new parameter samples and evaluate *d* until a fixed number of parameter samples are accepted. For each iteration, the fraction of accepted samples out of all drawn parameter samples is recorded as the acceptance rate *accR*. Next, the proposal distribution is updated using the samples accepted on the current iteration, and the new error threshold is set at the first quartile of the distance *d* for the accepted samples. The iterations continue until the acceptance rate reaches a specified value. The last set of accepted parameter samples is treated as an approximation for the posterior distribution (Methods 4.2). In the regular ABC algorithm, the error threshold is fixed and parameters are sampled from the prior distribution (same as in the first step of aABC). Updating the error threshold and proposal distribution in successive iterations of aABC allows for a more efficient fitting procedure especially when setting wide prior distributions [58]. We implemented this algorithm in the abcTau Python package, which includes different types of summary statistics and generative models that can be flexibly adjusted to account for various types of dynamics and noise in the biological data (Methods 4.3). To visualize the posterior distribution of a parameter (e.g., a timescale), we marginalize the obtained multivariate posterior distribution over all other parameters of the generative model.

We tested our method on the same synthetic data from the processes with known groundtruth timescales that we used to demonstrate the bias of direct fitting (Fig. 3 cf. Fig. 1). In these examples, we use the autocorrelation as the summary statistic and compute distances in linear scale. For all three processes, the shape of the sample autocorrelation of the observed data is accurately reproduced by the autocorrelation of synthetic data generated using the maximum *a posteriori* (MAP) estimate of parameters from the joint multivariate posterior distribution (Fig. 3, left). The posterior distributions inferred by the aABC algorithm include the ground-truth timescales (Fig. 3, middle). The variance of posterior distributions quantifies the uncertainty of estimates. In our simulations, the number of trials controls the signal to noise ratio in sample autocorrelation, and consequently the width of the posterior distributions (Supplementary Fig. 8).

**Fig. 3.**
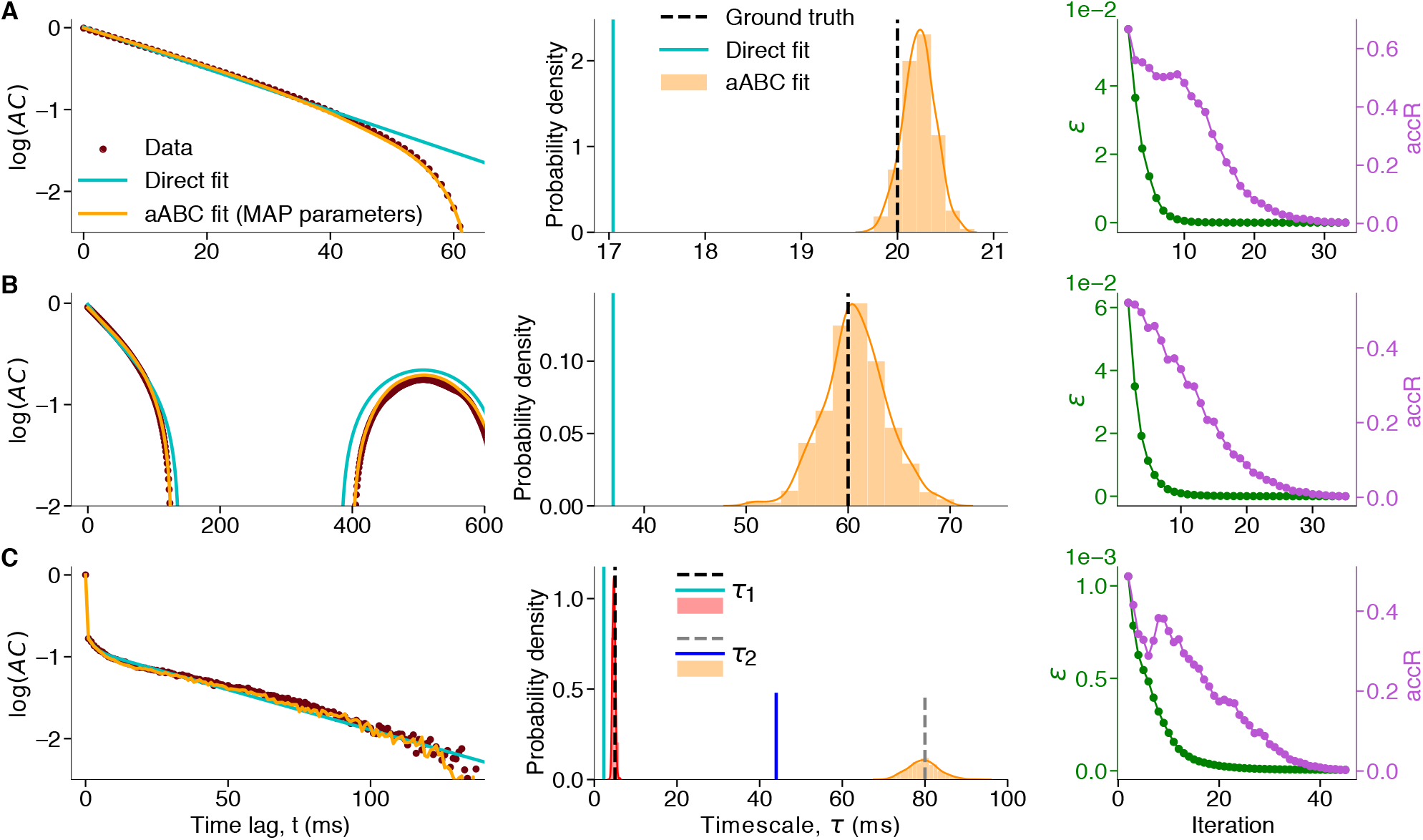
The aABC algorithm accurately recovers the ground-truth timescales and quantifies the estimation uncertainty. The same synthetic data as in Fig. 1 for *T* = 1 s. (**a**) Data are generated from an OU process with the timescale *τ* = 20 ms. **Left:** The shape of the data autocorrelation (brown) is accurately reproduced by the autocorrelation of synthetic data from the generative model with the MAP estimate parameters (orange, *τ*_MAP_ = 20.2 ms), but it cannot be captured by the direct exponential fit (cyan). **Middle:** The marginal posterior distributions (histograms overlaid with Gaussian kernel smoothing) include the ground-truth timescales, while direct exponential fits (cyan) underestimate the ground-truth timescales. The width of the posteriors indicates the estimation uncertainty. **Right:** The convergence of the aABC algorithm is defined based on the acceptance rate *accR* (purple), which decreases together with the error threshold *ε* (green) over the iterations of the algorithm. Data are plotted from the second iteration. Initial error threshold for all fits was set to 1. **(b)** Same as a for data from a linear mixture of an OU process with the timescale *τ* = 20 ms and an oscillation with frequency *f* = 2 Hz (Eq. 5). *τ*_MAP_ = 60.5 ms, **(c)** Same as a for data from an inhomogeneous Poisson process with two timescales *τ*_1_ = 5 ms and *τ*_2_ = 80 ms. *τ*_1,MAP_ = 4.7 ms, *τ*_2,MAP_ = 80 ms. See Methods 4.5 for other parameters.

Our method can also estimate timescales by fitting the sample PSD in the frequency domain. The aABC method recovers the ground-truth timescales by fitting the whole shape of the PSD without the need to tune the fitted frequency range, even in the presence of multiple timescales and additional noise (e.g., spiking noise) (Supplementary Fig. 9) or multiple oscillatory components (Supplementary Fig. 10). Moreover, aABC method can uncover slow oscillations in signals that do not exhibit clear peaks in PSD due to the short duration of time-series and the low frequency resolution (Supplementary Fig. 9C). The aABC method can be used in combination with any method for computing the PSD (e.g., any window functions for removing the spectral leakage), since the exact same method applied to synthetic data would alter its sample PSD in the same way.

Different summary statistics and fitting ranges may be preferred depending on the application. For example, autocorrelations allow for using additional correction methods such as jittering [46], whereas PSD estimation can be improved with filtering or multitapers [67]. Further, selecting a smaller *t_m_* when fitting autocorrelations prevents overfitting to noise in the autocorrelation tail. Distances between summary statistics can also be computed on the logarithmic scale (e.g., log(AC) or log(PSD)). The choice of summary statistic (metric and fitting range) can influence the shape of the approximated posterior (e.g., posterior width), but the posteriors peak close to the ground-truth timescales as long as the same summary statistic is used for observed and synthetic data (Supplementary Fig. 11). The advantage of such behavior is particularly visible when changing the range over which we compute the summary statistics (e.g., minimum and maximum frequency in PSD). The direct fitting is sensitive to the selected range and can lead to wrong estimates of timescales, whereas aABC recovers correct timescales for all ranges (Supplementary Fig. 6).

In the aABC algorithm, we set a multivariate uniform prior distribution over the parameters of the generative model. The ranges of uniform prior should be broad enough to include the ground-truth values. Selecting broader priors does not affect the shape of posterior distributions obtained with the aABC and only slows down the fitting procedure (Supplementary Fig. 12). Hence, we can set wide prior distributions when a reasonable range of parameters is unknown. Timescales estimated from the direct exponential fits can be used as a potential lower bound for the priors.

We evaluated the reliability of the aABC method on a wide range of timescales and trial durations and compared the results to direct fitting (Supplementary Fig. 13). The estimation error of direct fitting increases when trial durations become short relative to the timescale, whereas the aABC method always returns reliable estimates. However, the estimation of the full posterior distribution comes with a price of higher computational costs (Supplementary Table 1) compared to point estimates. Thus, the direct fit of the sample autocorrelation may be preferred when the long time-series data is available so that statistical bias does not corrupt the results. To empirically verify whether this is the case, we implemented a pre-processing algorithm in our Python package that uses parametric bootstrapping to return an approximate error bound of the direct fit estimates (Methods 4.3.1, Supplementary Fig. 14).

Fitting generative models with the aABC method provides a principled framework for estimating timescales that can be used with different metrics in time or frequency domain. Furthermore, joint posterior distribution of the inferred parameters allows us to examine correlations between different parameters, find manifolds of possible solutions, and identify a potential degeneracy in the parameter space (Supplementary Fig. 15).

### 2.3 Estimating the timescale of activity in a branching network model

So far, we demonstrated that our aABC method accurately recovers the ground-truth timescales within the same model class, i.e. when the generative model and the process that produced the observed data are the same. However, the timescale inference based on OU processes is broadly applicable when the mechanism that generated the exponential decay of autocorrelation in the data is not an OU process.

Here, we tested our inference method on discrete-time data from an integer-valued autoregressive model with a known ground-truth autocorrelation function. Specifically, we applied our method to estimate the timescale of the global activity in a branching network model [68–70]. This model is often used to study the operating regime of dynamics in neuronal networks, e.g., how close the dynamics is to the critical state. A branching network consists of *k* interconnected binary neurons, each described by the state variable *x_i_* ∈ {0, 1}, where *x_i_* = 1 indicates that neuron i is active, and 0 that it is silent (Fig. 4A). We considered a fully-connected network. Each active neuron can activate other neurons with the probability *p* = *m/k* and then, if not activated by other neurons, it becomes inactive again in the next time-step. Additionally, at every time-step, each neuron can be activated with a probability *h* by an external input. For a small input strength *h*, the state of the network’s dynamics is governed by a branching parameter *m* (*m* = 1 corresponds to the critical state). The autocorrelation function of the global activity *A*(*t′*) = Σ_*i*_ *x_i_*(*t′*) in this network is known analytically *AC*(*t_j_*) = exp(*t_j_*ln(*m*)) [25]. Thus the ground-truth timescale of this activity is given by *τ* = – 1/ ln(*m*).

**Fig. 4.**
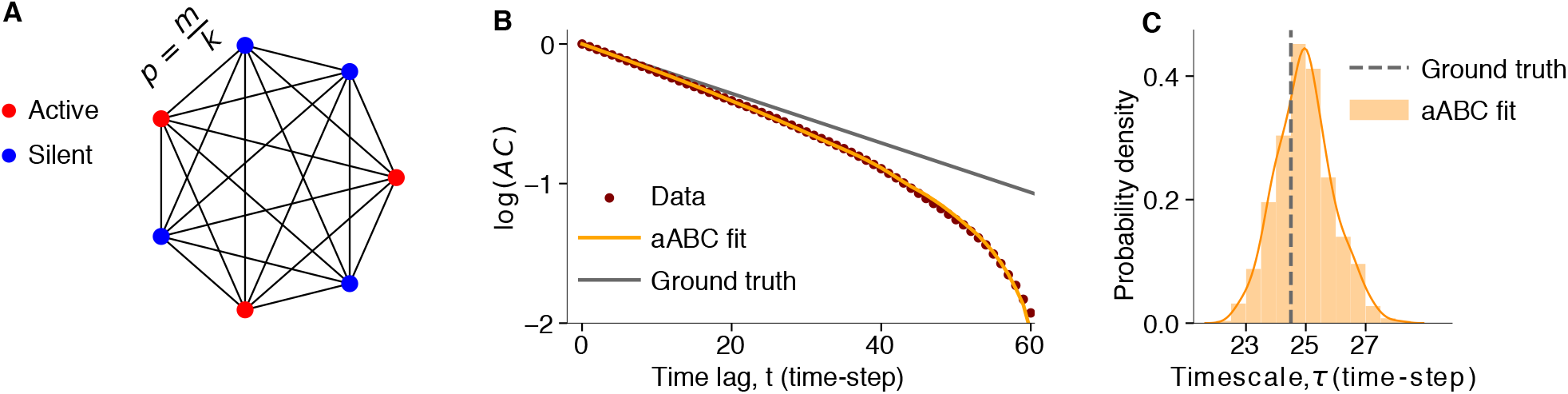
The aABC inference with a generative model based on the OU process recovers the ground-truth timescale of activity in a branching network. (**a**) A schematic of a fully connected branching network with binary neurons. Branching parameter (*m* = 0.96) and number of neurons (*k* = 10^4^) define the probability of activity propagation. Each neuron receives an external input with probability *h* = 10^-3^. (**b**) The shape of the sample autocorrelation of simulated activity in the branching network (brown) deviates from the ground truth (gray), but is accurately reproduced by the autocorrelation of synthetic data generated from a one-timescale OU process with the MAP estimate timescale from the aABC method. *τ*_MAP_ = 24.9. The data contained 100 trials of 500 timesteps. (**c**) The aABC posterior distribution includes the ground-truth timescale. *τ*_ground truth_ = 24.5, *τ*_aABC_ = 24.9 ± 0.9 (mean ± std). Fitting parameters are provided in Methods 4.5.

We simulated the binary network model to generate the time-series data of the global activity. We then used aABC to fit the sample autocorrelation of these data with a one-timescale OU process as the generative model. The inferred posterior distribution is centered on the theoretically predicted timescale of the branching network, and the MAP estimate parameter accurately reproduces the shape of the sample autocorrelation of the network activity (Fig. 4B,C). These results show that our framework can be used to estimate timescales in diverse types of data with different underlying mechanisms.

### 2.4 Model selection with Approximate Bayesian Computations

For experimental data, the correct generative model is usually unknown. For example, it may not be obvious *a priori* how many timescales should the generative model contain. Several alternative hypotheses may be plausible, and we need a procedure to select which one is more likely to explain the observed data. Assuming that the autocorrelation function or PSD is a sufficient summary statistic for estimating timescales, we can use aABC to approximate the Bayes factor for selecting between alternative models [71–73]. Model selection based on ABC can produce inconsistent results when the summary statistic is insufficient [74], but whether this is likely the case can be verified empirically [73]. Specifically, the summary statistic can be used for model selection with ABC if their mathematical expectation is significantly different for the two models.

Based on this empirical procedure [73], we developed a method for selecting between two alternative models *M*_1_ and *M*_2_ using their aABC fits. Models are compared using a goodness of fit measure that describes how well each model fits the data. The goodness of fit can be measured by the distance *d* between the summary statistic (e.g., autocorrelation or PSD) of synthetic and observed data (i.e. residual errors, Eq. 9). For fair comparison, the same summary-statistics and fitting range should be used for fitting both models. Since *d* is a noisy measure because of the finite sample size and uncertainty in the model parameters, we compare the distributions of distances generated by two models with parameters sampled from their posterior distributions. To approximate the distributions of distances, we generate multiple samples of synthetic data from each model with parameters drawn from its posterior distribution and compute the distance *d* for each sample. If the distributions of distances are significantly different (i.e. expectations of the summary statistic for two models are significantly different [73]), then we continue with the model selection, otherwise the summary statistic is insufficient to distinguish these models.

For selecting between *M*_1_ and *M*_2_, we use the distributions of distances to estimate the Bayes factor, which is the ratio of marginal likelihoods of the two models and accounts for the model complexity [75]. Assuming both models are *a priori* equally probable (*p*(*M*_1_) = *p*(*M*_2_)), the Bayes factor can be approximated using the models’ acceptance rates for a specific error threshold *ε* [71, 74, 76]

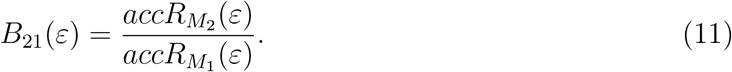

*B*_21_(*ε*) > 1 indicates that the model *M*_2_ is more likely to explain the observed data and vice versa. To eliminate the dependence on a specific error threshold, we compute the acceptance rates and the Bayes factor with a varying error threshold. For each error threshold *ε*, the acceptance rate is given by the cumulative distribution function of the distances CDF_*M_i_*_ (*ε*) = *p_M_i__* (*d* < *ε*) = *accR_M_i__* (*ε*) (*i* = 1, 2). Hence, the ratio between CDFs of two models gives the value of the Bayes factor for every error threshold *B*_21_ (*ε*) = CDF_*M*_2__ (*ε*)/CDF_*M*_1__ (*ε*). We select the model *M*_2_ if *B*_21_(*ε*) > 1, i.e. if CDF_*M*_2__ (*ε*) > CDF_*M*_1__ (*ε*) for all *ε*, and vice versa.

We evaluated our model selection method using synthetic data from three example processes with known ground truth, so that the correct number of timescales is known. Specifically, we used an OU process with a single timescale (Fig. 5A) and two different examples of an inhomogeneous Poisson process with two timescales. In the first example of an inhomogeneous Poisson process, the ground-truth timescales were well separated, so that the shape of the data autocorrelation suggested that the underlying process had multiple timescales (Fig. 5D). In the second example of an inhomogeneous Poisson process, we chose the ground-truth timescales to be more similar, so that a simple visual inspection of the data autocorrelation could not clearly suggest the number of timescales in the underlying process (Fig. 5G). For all three example processes, we fitted the data with one-timescale (*M*_1_) and two-timescale (*M*_2_) generative models using aABC and selected between these models by computing the Bayes factors. The one- and two-timescale models were based on a single OU process or a linear mixture of two OU processes, respectively. For the data from inhomogeneous Poisson processes, the generative model also incorporated an inhomogeneous Poisson noise.

**Fig. 5.**
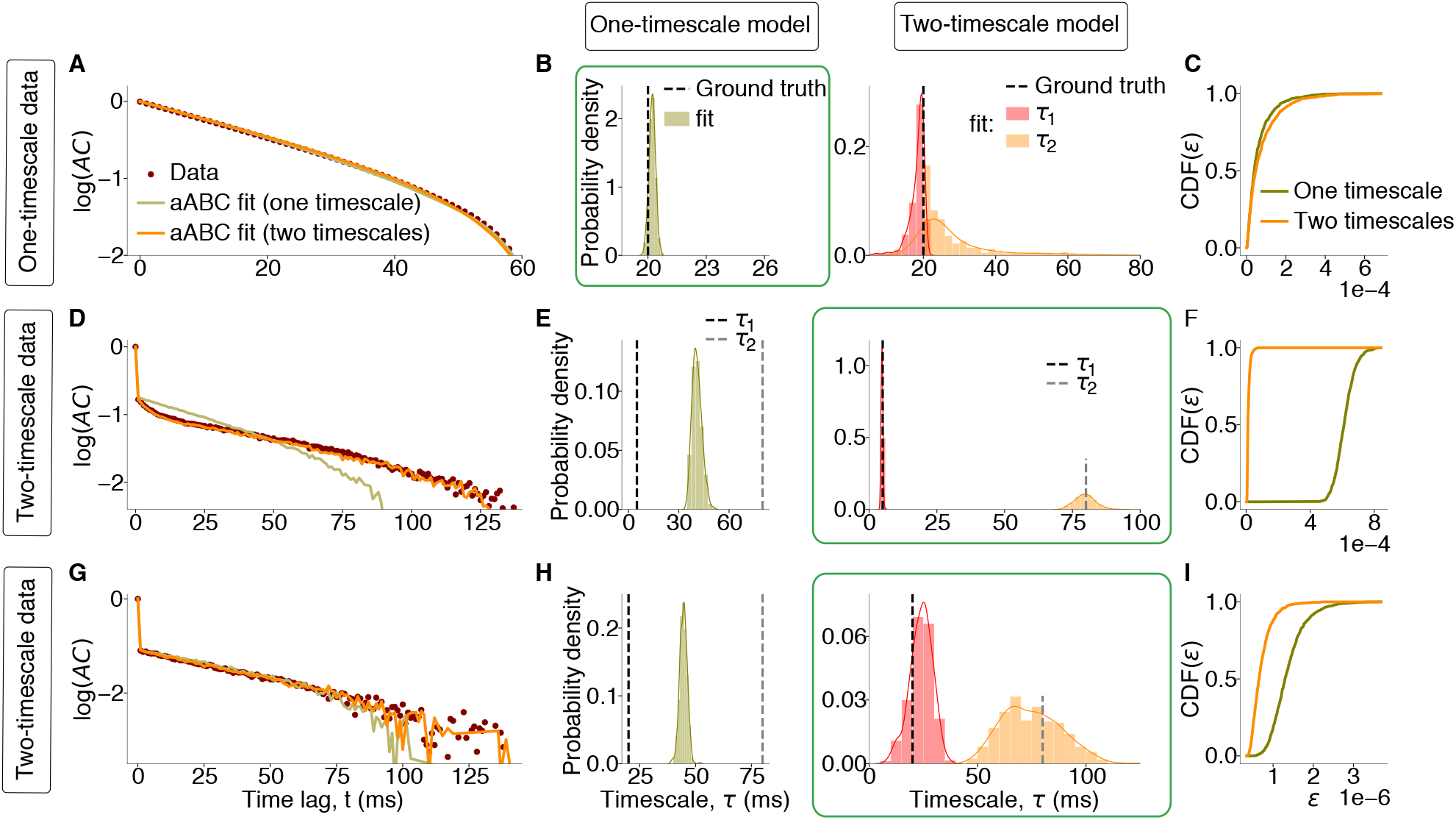
Model selection with ABC. (**a-c**) Data are generated from a one-timescale OU process with *τ* = 20 ms. (**a**) The data autocorrelation (brown) is fitted with a one-timescale (olive, *M*_1_) and two-timescale (orange, *M*_2_) generative models (sample autocorrelations for MAP estimate parameters are shown). (**b**) Marginal posterior distribution of the timescale estimated using the one-timescale model includes the ground truth and has small variance (**left**). Marginal posterior distributions of two timescales estimated using the two-timescale model heavily overlap and have large variance (**right**). (**c**) Cumulative distributions of distances *d* for two models, CDF*_M_i__*(*ε*). Since CDF_*M*_2__ (*ε*) < CDF_*M*_1__ (*ε*) for all *ε*, the one-timescale model is selected (green box in B). (**d-f**) Data are generated from an inhomogeneous Poisson process with two timescales: *τ*_1_ = 5 ms, *τ*_2_ = 80 ms. (**d**) Same format as A. (**e**) Marginal posterior distributions for two timescales estimated by the two-timescale model include the ground truth (**right**). Marginal posterior distributions for the timescale estimated by the one-timescale model falls in between the ground-truth timescales (**left**). (**f**) Cumulative distributions of distances *d* for two models. Since CDF_*M*_2__ (*ε*) > CDF_*M*_1__ (*ε*) for all *ε*, the two-timescale model is selected (green box in E). (**g-i**) Same as d-f, but for *τ*_1_ = 20 ms, *τ*_2_ = 80 ms. For all fits, CDF*_M_i__* (*ε*) are computed from 1000 samples. Simulation and fitting parameters are provided in 4.5.

For the example OU process with a single timescale, the one- and two-timescale models fitted the shape of the data autocorrelation almost equally well (Fig. 5A). The marginal posterior distributions of two timescales estimated by the two-timescale model heavily overlap around their peak values (Fig. 5B), which indicates that the one-timescale model possibly better describes the data. For the two-timescale model we enforce the ordering *τ*_1_ < *τ*_2_, which generates the difference between their distributions. To select between two models, we compare the cumulative distributions of distances CDF_*M_i_*_ (*ε*) (Fig. 5C). Although the two-timescale model has more parameters, it has significantly larger distances than the one-timescale model (Wilcoxon rank-sum test, *P* = 0.002, mean *d*_*M*_1__ = 6 × 10^-5^, mean *d*_*M*_2__ = 8 × 10^-5^). The two-timescale model has a larger average distance because its posterior distribution has larger variance (larger uncertainty), which leads to a higher possibility to sample a combination of parameters with a larger distance. Since CDF_*M*_2__ (*ε*) < CDF_*M*_1__ (*ε*) (i.e. *B*_21_(*ε*) < 1), the one-timescale model is preferred over the two-timescale model, in agreement with the ground-truth generative process.

For both example inhomogeneous Poisson processes with two timescales, the shape of the data autocorrelation is better matched by the two-timescale than by the one-timescale model (the difference is subtle for the second example, Fig. 5D,G). The marginal posterior distributions of two timescales estimated by the two-timescale model are clearly separated and include the ground-truth values, whereas the timescale estimated by the one-timescale model is in between the two ground-truth values (Fig. 5E,H). The two-timescale model has significantly smaller distances (Wilcoxon rank-sum test, Fig. 5F: *P* < 10^-10^, mean *d*_*M*_1__ = 6 × 10^-4^, mean *d*_*M*_2__ = 1.5 × 10^-5^; Fig. 5I: *P* < 10^-10^, mean *d*_*M*_1__ = 10^-6^, mean *d*_*M*_2__ = 7 × 10^-7^). Since CDF_*M*_2__(*ε*) > CDF_*M*_1__ (*ε*) (i.e. *B*_21*d*_(*ε*) > 1) for all error thresholds, the two-timescale model provides a better description of the data for both examples, in agreement with the ground truth. Thus, our method selects the correct generative model even for a challenging case where the shape of the data autocorrelation does not suggest the existence of multiple timescales. Our method can be used to discriminate between a broad class of models, e.g., slow periodic (e.g., slow oscillation) versus aperiodic (e.g., slow exponential decay) components in the time-series (Supplementary Fig. 16).

### 2.5 Estimating timescales of ongoing neural activity

To illustrate an application of our computational framework to experimental data, we inferred the timescales of ongoing spiking activity in the primate visual cortex. The spiking activity was recorded from the visual area V4 with a 16-channels micro-electrode array [77]. During recordings, a monkey was fixating a central dot on a blank screen for 3 s on each trial. To estimate the timescales, we pooled the spiking activity recorded across all channels and computed the spike-count autocorrelation of the pooled activity with a bin-size of 1 ms.

Previously, the autocorrelation of neural activity in several brain regions was modeled as an exponential decay with a single timescale [3]. To determine whether a single timescale is sufficient to describe the temporal dynamics of neural activity in our data, we compared the one-timescale (*M*_1_) and two-timescale (*M*_2_) models and selected the model that better described the data. As a generative model for neural spike-counts, we used a doubly-stochastic process [64,65], where spike-counts are generated from an instantaneous firing rate modelled as one OU process (*M*_1_) or a mixture of two OU processes (*M*_2_). To account for the non-Poisson statistics of the spike-generation process, we sampled spike-counts from a gamma distribution [78] (Methods 4.1.3). We fitted both models with the aABC algorithm and selected between the models using Bayes factors (Fig. 6A-C). The two-timescale model provided a better description for the data, since it had smaller distances and CDF_*M*_2__ (*ε*) > CDF_*M*_1__ (*ε*) for all error thresholds (Fig. 6C, Wilcoxon rank-sum test, *P* < 10^-10^, mean *d*_*M*_1__ = 8 × 10^-4^, mean *d*_*M*_2__ = 2 × 10^-4^).

**Fig. 6.**
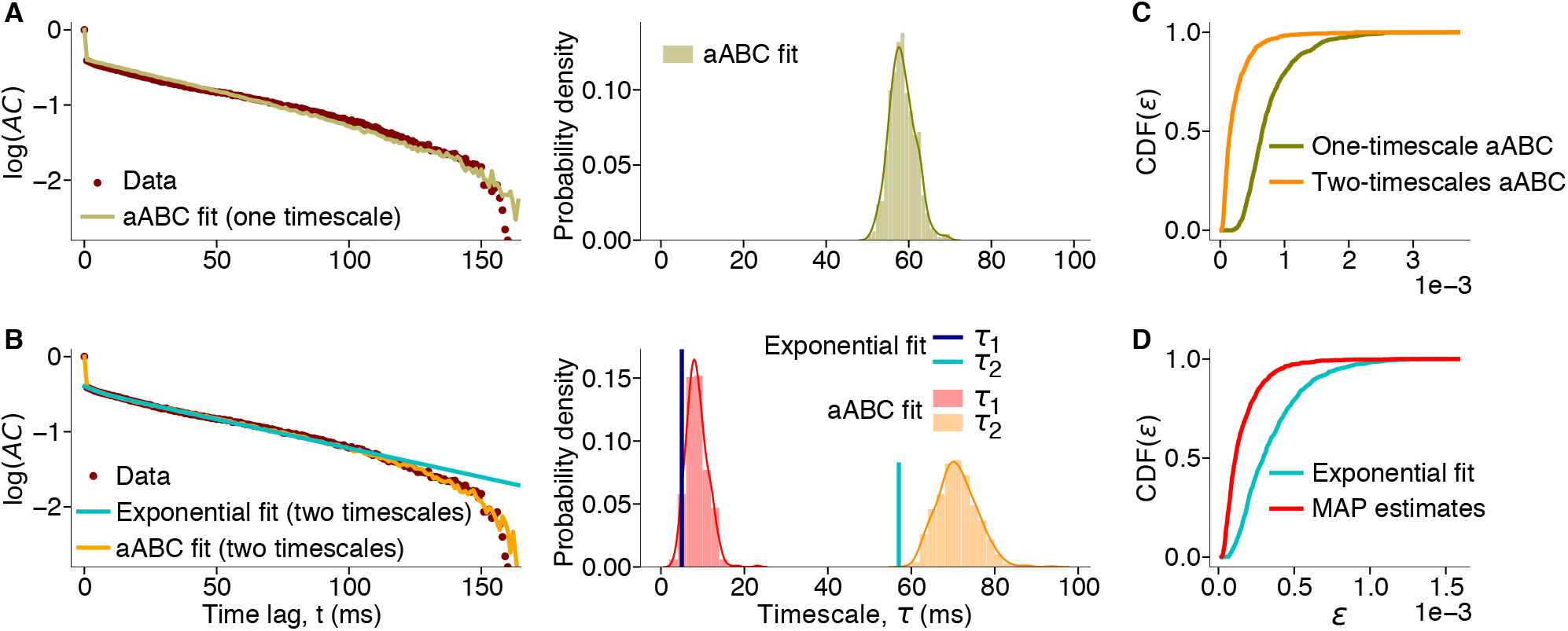
Estimating timescales of ongoing neural activity and comparing hypotheses about their number with aABC. (**A**) **Left:** The autocorrelation of neural spike-counts (brown) is fitted with a one-timescale doubly-stochastic model using aABC (olive, sample autocorrelation for MAP estimate parameters). **Right:** Posterior distribution of the timescale. *τ*_MAP_ = 58 ms. (**B**) **Left:** Same data as in A is fitted directly with a double exponential function (cyan) and with a two-timescale doubly-stochastic model using aABC (orange, sample autocorrelation for MAP estimate parameters). **Right:** Timescales estimated by the direct exponential fit (*τ*_1_ - blue, *τ*_2_ -cyan) and marginal posterior distributions of timescales inferred with aABC (*τ*_1_ - red, *τ*_2_ - orange). *τ*_1,exp_ = 5 ms, *τ*_2,exp_ = 57 ms, *τ*_1,MAP_ = 8 ms, *τ*_2,MAP_ = 70 ms. (**C**) Cumulative distribution of distances *d* for one-timescale (*M*_1_) and two-timescale (*M*_2_) models. Since CDF_*M*_2__ (*ε*) > CDF_*M*_1__ (*ε*) for all *ε*, the two-timescale model is selected. (**D**) Cumulative distributions of distances *d* between the autocorrelation of neural data and synthetic data from the two-timescale doubly-stochastic model with parameters either from the direct fit (cyan) or MAP estimate with aABC (red). The MAP parameters have smaller distances, i.e. describe the autocorrelation of neural data more accurately than the direct fit. Fitting parameters are provided in Methods 4.5.

We further compared our method with a direct exponential fit of the sample autocorrelation which is usually employed to infer the timescales of neural activity [3, 23, 25, 29–31]. We fitted the sample autocorrelation with a double exponential function

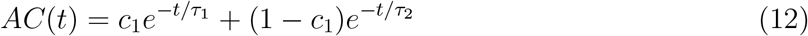

up to the same *t_m_* = 150 ms as in the aABC method and compared the result with the two-timescale aABC fit (Fig. 6B). Similar to what we observed with synthetic data, the direct exponential fit produced timescales that were systematically smaller than the MAP estimates with the aABC fit. Including all the time lags in exponential fitting results in even larger bias. Since the ground-truth timescales are not available for biological data, we used a sampling procedure to evaluate whether the data is better described by the timescales from the direct fit or from the MAP estimates with the aABC fit. We generated multiple samples of synthetic data using the two-timescale doubly-stochastic generative model with the parameters from either the direct fit or MAP estimates from aABC. For each sample, we measured the distance *d* between the autocorrelation of synthetic and neural data to obtain the distribution of distances for both types of fits (Fig. 6D). The distances were significantly smaller for synthetic data generated with the parameters from the MAP estimate with aABC than from the direct exponential fit (Wilcoxon rank-sum test, *P* < 10^-10^, mean distance of MAP parameters 10^-4^, mean distance of exponential fit parameters 3× 10^-4^). Our method performs better because it accounts for the bias in the autocorrelation shape which is ignored by the direct fit. These results suggest that our method estimates the timescales of neural activity more accurately than a direct exponential fit and, moreover, allows for comparing alternative hypotheses about the underlying dynamics.

## 3 Discussion

We demonstrated that direct fitting the shape of sample autocorrelation or PSD often fails to recover the correct timescales due to a statistical bias arising from finite sample size. Since the bias depends on the duration of time-series (Fig. 1, left), comparing timescales estimated from direct fits of autocorrelations or PSDs of experimental data with different duration can lead to misleading interpretations.

To overcome this problem, we developed a flexible computational framework based on adaptive Approximate Bayesian Computations that is broadly applicable to different types of data. Our framework fits the sample autocorrelation or PSD of the data with a generative model which is based on a mixture of Ornstein-Uhlenbeck processes (one for each estimated timescale) and can incorporate additional temporal structure and noise. Our method infers a posterior distribution of timescales consistent with the observed data. The width of the posterior distribution depends on the noise and amount of data and quantifies the estimation uncertainty, e.g., it provides a confidence interval for estimated timescales. The posterior distributions can be used for model selection to compare alternative hypotheses about the dynamics of the underlying process. This framework is suitable for comparing timescales between datasets with different durations and between processes that exhibit different temporal dynamics (e.g., oscillations). Our approach is not sensitive to the specific method for computing autocorrelations or PSDs and does not require fine-tuning of the fitting ranges, since the same method applies to observed and synthetic data.

While previous work [56] attributed errors in the estimated timescales to fitting noise in the tail of autocorrelation, we find that the main reason is the statistical bias in the sample autocorrelation. This bias arises primarily due to the deviation of the sample mean from the ground-truth mean. If the ground-truth mean of the process is known, then using the true mean instead of the sample mean for computing the autocorrelation largely eliminates the bias. In this case, direct fitting can produce satisfactory estimates, but our method additionally provides the measure of estimation uncertainty. When the true mean is unknown but it is known that it is the same across all trials, it is beneficial to estimate a single sample mean from the whole dataset instead of estimating the mean for trials individually. However, this assumption does not always hold. For example, the mean of spiking neural activity can change across trials because of changes in animal’s behavioral state. If the assumption of a constant mean is violated in the data, estimating the sample mean from the whole dataset leads to strong distortions of the autocorrelation shape introducing additional slow timescales [79].

We validated our method using data from synthetic processes, where the exponential decay rate of autocorrelation is known analytically by design. We further illustrated the application of our method to spiking neural activity data, where the underlying ground-truth process is unknown. In this case, the posterior distributions estimated by our method can be used to approximate the Bayes factor for selecting between different generative models that represent alternative hypotheses about neural dynamics. Since the Bayes factor takes the model complexity into account, it can be used for comparing models with different number of parameters, e.g., to infer the number of timescales in the data.

Here, we focused on exponentially decaying autocorrelation shapes (i.e. Lorentzian PSD shape with the 1/*f* power-law exponent of 2) since the definition of timescales based on the exponential decay rates of the autocorrelation is widely used in the literature [3,25,29–40] and has a clear interpretation as a timescale of a generative dynamical system. While it is true that many types of data exhibit PSD with 1/*f* exponents deviating from 2 [42,80], there is no universally accepted answer to what would be the definition of timescale for these types of data and what is the nature of processes generating this behavior in PSD. Indeed, a prominent hypothesis is that the 1/*f* PSD arises from a mixture of processes with exponential autocorrelations and different timescales [22, 42, 80, 81] that can be modeled as a mixture of OU processes. For example, in neuroscience, a combination of excitatory (AMPA) and inhibitory (GABA) synaptic currents with distinct timescales was suggested as a potential mechanism for creating different 1/*f* exponents [80]. However, sometimes we are interested in separating a process with a well-defined timescale from 1/*f* background. To handle such cases, we introduced an augmented generative model (Methods 4.1.4) which can be used to estimate timescales in the presence of a background process with an arbitrary shape of PSD (e.g., 1/*f* exponents other than 2). This generative model is agnostic to the nature of the process generating the background activity and directly models the desired PSD shape. We can use this model to simultaneously estimate the 1/*f* exponent and exponential decay timescales (Supplementary Fig. 17).

The general framework of inferring timescales with aABC based on OU processes can be adapted to various types of data, different generative models and summary statistics using our Python package. We implemented a set of generative models, summary statistic computation methods and distance functions in the abcTau package. Moreover, the modular implementation of our Python package allows users to easily incorporate additional types of dynamics and non-stationarities into customized generative models, or use other types of summary statistic which can be added directly to the package. Thus, the package can be extended to a wide range of applications in different fields including neuroscience, cellular biology, epidemiology, and physics. Our framework provides a principled method for estimating timescales using summary statistics in time or frequency domain. This approach is particularly favorable for data organized in short trials or trials of different durations, when standard direct fitting is unreliable, and it allows for comparing timescales between datasets with different durations and data amount.

## 4 Methods

### 4.1 Generative models

We used several generative models based on a linear mixture of OU processes—one for each estimated timescale—sometimes augmented with additional temporal structure (e.g., oscillations) and noise.

#### 4.1.1 Ornstein–Uhlenbeck process with multiple timescales

We define an OU process with multiple timescales *A*(*t′*) as a linear mixture of OU processes *A_k_*(*t′*) with timescales *τ_k_*, *k* ∈ {1,…, *n*}, zero mean and unit variance:

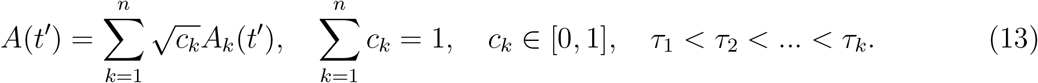

Here *n* is the number of timescales in the mixture and *c_k_* are the mixing coefficients. We simulate each OU process *A_k_* by iterating its time-discrete version using the Euler scheme [61]

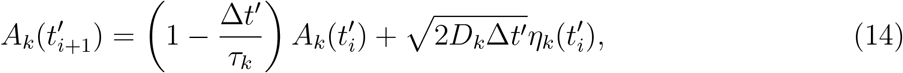

where 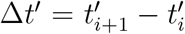 is the discretization time-step and 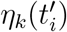 is a random number generated from a normal distribution. We set the unit variance for each OU process Var(*A_k_*) = *D_k_τ_k_* = 1 by fixing *D_k_* = 1/*τ_k_*. The parameter-vector *θ* for a linear mixture of *n* OU processes consists of 2*n* – 1 values: *n* timescales *τ_k_* and *n* – 1 coefficients *c_k_* (see Eq. 13).

We match the mean and variance of the multi-timescale OU process to the sample mean 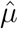 and sample variance 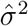 of the observed data using a linear transformation:

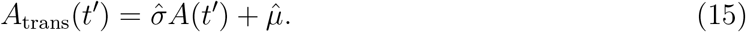

We use the process *A*trans(*t′*) as a generative model for data fitting and hypothesis testing (Methods 4.2).

#### 4.1.2 Multi-timescale Ornstein–Uhlenbeck process with an oscillation

To obtain a generative model with an oscillation, we add to a multi-timescale OU process (Eq. 13) an oscillatory component with the weight *c*_*k*+1_:

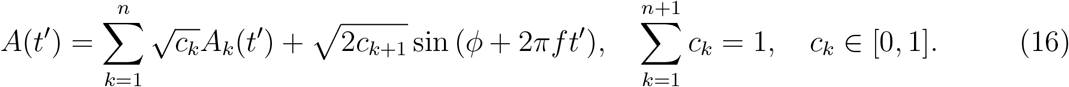

For the analysis in Fig. 1B and in Fig. 3B, we assumed that the frequency *f* is known and only fitted the weight *c*_*k*+1_, showing that the bias problem for the direct fit persists even if we know the correct frequency of oscillation. The frequency *f* can be fitted with aABC as an additional free parameter (Supplementary Fig. 9 and 10). For each trial, we draw the phase φ independently from a uniform distribution on [0, 2*π*]. We use the linear transformation in Eq. 15 to match the mean and variance of this generative process to the observed data.

#### 4.1.3 Doubly stochastic process with multiple timescales

The doubly stochastic process with multiple timescales is generated in two steps: first generating time-varying rate and then generating event-counts from this rate. To generate the time-varying rate, we scale, shift and rectify a multi-timescale OU process (Eq. 13) using the transformation

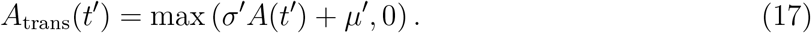

The resulting rate *A*_trans_(*t′*) is non-negative and for *μ′* ≫ *σ* it has the mean 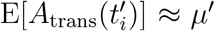 and variance 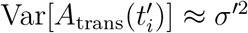. We then draw event-counts s for each time-bin 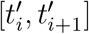 from an event-count distribution 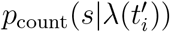, where 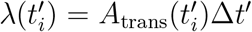 is the mean event-count and 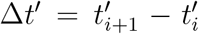 is the bin size (in our simulations Δ*t′* = 1 ms). A frequent choice of 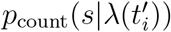 is a Poisson distribution

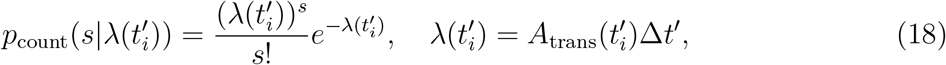

which results in an inhomogeneous Poisson process.

To match the mean and variance of the doubly stochastic process to the observed data, we need to estimate the mean rate *μ′* and both the variance of the rate *σ′*^2^ and the variance of the event-count distribution 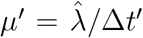. According to the law of total expectation, the mean rate 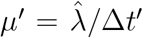, where 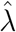 is the sample mean of the observed event-counts. According to the law of total variance [65], the total variance of event-counts *σ*^2^ arises from two contributions: the variance of the rate and the variance of the event-count distribution:

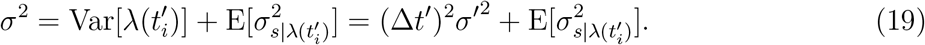

For the Poisson distribution, the variance of the event-count distribution is equal to its mean: 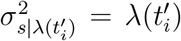. However, the condition of equal mean and variance does not always hold in experimental data [78]. Therefore, we also use other event-count distributions, in particular a Gaussian and gamma distribution. We define *α* as the variance over mean ratio of the eventcount distribution 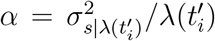. For the Poisson distribution, *α* = 1 always holds. For other distributions, we assume that *α* is constant (i.e. does not depend on the rate). With this assumption, the law of total variance Eq. 19 becomes

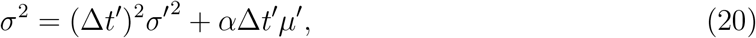

where 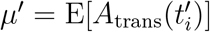 is the mean rate. From Eq. 20 we find the rate variance

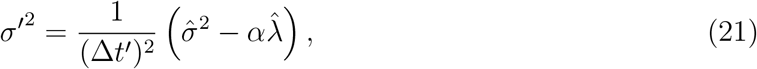

where 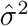 is the sample variance of event-counts in the observed data. We find that with the correct values of *μ′*, *σ′*^2^ and *α*, both the Gaussian and gamma distributions of event-counts produce comparable estimates of timescales (Supplementary Fig. 18).

To calculate *σ′*^2^ with Eq. 21, we first need to estimate α from the observed data. We estimate α from the drop of autocorrelation of event-counts between the time-lags *t*_0_ and *t*_1_. Since eventcounts in different time-bins are drawn independently, this drop mainly reflects the difference between the total variance of event-counts and variance of the rate (Eq. 19, we neglect a small decrease of the rate autocorrelation between *t*_0_ and *t*_1_) and does not depend on timescales. Thus, we find α with a grid search that minimizes the distance between the autocorrelation at *t*_1_ of the observed and synthetic data from the generative model with fixed timescales. Alternatively, α can be fitted together with all other parameters of the generative model using aABC. We find that since α is almost independent from other parameters, aABC finds the correct value of α first and then fits the rest of parameters. The MAP estimate of α converges to the same value as estimated by the grid search, but aABC requires more iterations to get posterior distributions for estimated timescales with a similar variance (Supplementary Fig. 19). Therefore, gridsearch method is preferred when moderate errors in α are acceptable and approximate range of ground-truth timescales are known, but for more accurate results it is better to fit the α by aABC together with other parameters.

To estimate the timescales of the two-timescale inhomogeneous Poisson process with exponential fits (Fig. 1C), we assumed that the mean and variance of the event-counts are known and only estimated *τ*_1_, *τ*_2_ and the coefficient *c*_1_ by fitting Eq. 7 to the sample autocorrelation starting from the lag *t*_1_.

#### 4.1.4 Modelling background processes with an arbitrary PSD shape

We can generate random time-series with any desired shape of the PSD. First, we convert the power spectrum PSD(f) to amplitudes 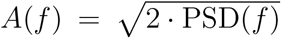. Then, we draw random phases *ϕ*(*f*) from a uniform distribution on [0, 2*π*] and construct a frequency domain signal *Z*(*f*) = *A*(*f*)*e*^*iϕ*(*f*)^. We transform this signal to the time domain by applying the inverse Fast Fourier Transform (iFFT) and z-score the time-series to obtain zero mean and unit variance.

For example, a common class of background processes exhibit power-law decay in PSD [22,42, 80, 81]. We model this PSD shape as PSD(*f*) = *f^-χ^*, where *f*_min_ < *f* < *f*_max_ is the frequency range and *χ* is the power-law exponent. We can set the lower frequency cut-off at *f*_min_ = 1 Hz and the upper frequency cut-off *f*_max_ is defined by the Nyquist frequency (i.e. half of the desired sampling rate). We can combine this process with time-series generated from an OU process with the timescale *τ* to obtain a PSD shape with both 1/*f* and Lorentzian components (Supplementary Fig. 17). We sum the two time-series with the coefficients *c* and 1 – *c* and rescale using the linear transformation Eq. 15 to match the mean and variance to the observed data. This generative process can be used when the 1/*f* exponent in the data PSD deviates from 2 (i.e. Lorentzian shape). This method can be applied to generate background processes with arbitrary desired PSD shapes and estimate the relevant parameters.

### 4.2 Optimizing generative model parameters with adaptive Approximate Bayesian Computations

We optimize parameters of generative models with adaptive Approximate Bayesian Computations (aABC) following the algorithm from Ref. [58]. aABC is an iterative algorithm to approximate the multivariate posterior distribution of model parameters. It uses population Monte-Carlo sampling to minimize the distance between the summary statistic of the observed and synthetic data from the generative model. We use sample autocorrelation or PSD as the summary statistic and define the distance *d* as the mean squared deviation between the observed and synthetic data summary statistics (Eq. 9, Eq. 10).

On the first iteration of the algorithm, the parameters of the generative model 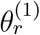 are drawn from the prior distribution. We use a multidimensional uniform prior distribution *π*(*θ*) over fitted parameters (e.g., timescales and their weights). The domain of prior distribution for the timescales is chosen to include a broad range below and above the timescales estimated by the direct exponential fits of data autocorrelation (Table 2). For the weights of timescales *c_k_*, we use uniform prior distributions on [0, 1]. The model with parameters 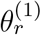 is used to generate synthetic time-series *A*(*t′*) with the same duration and number of trials as in the observed data. Next, we compute the distance *d* between the summary statistics of the synthetic and observed data. If *d* is smaller than the error threshold *ε* (initially set to 1), the parameters are accepted and added to the multivariate posterior distribution. Each iteration of the algorithm is repeated until 500 parameters samples are accepted.

**Table 1.**
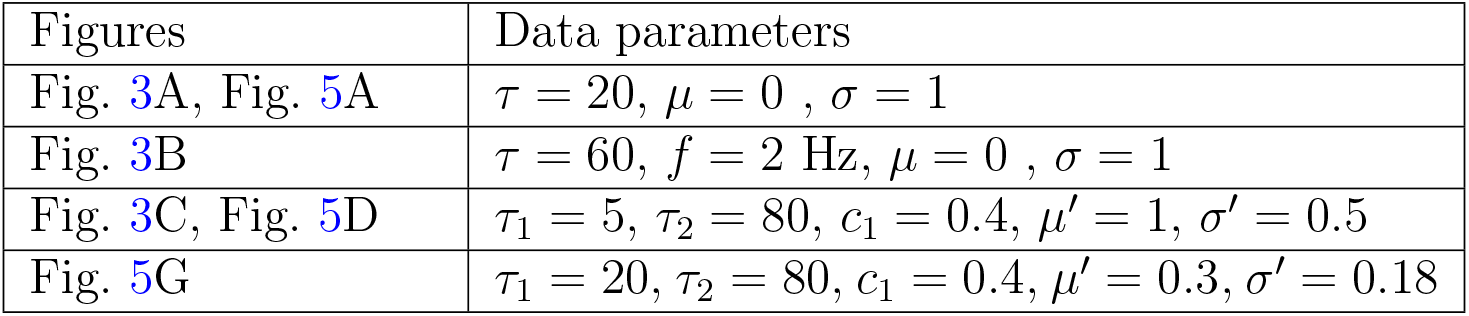
Simulation parameters for data autocorrelations.

**Table 2.** aABC fit parameters. *U*[a, b] denotes a uniform distribution on the interval [a, b].

| Figures | Priors | $t_m$ |
| --- | --- | --- |
| Fig. 3A, Fig. 4, Fig. 5B | $\tau, \tau_1, \tau_2 : U[0, 60]$ | 50 |
| Fig. 3B | $\tau : U[0, 120]$ | 100 |
| Fig. 3C, Fig. 5E (Right) | $\tau_1 : U[0, 60], \tau_2 : U[20, 140]$ | 110 |
| Fig. 5E (Left) | $\tau : U[0, 140]$ | 110 |
| Fig. 5H (Right) | $\tau_1 : U[0, 60], \tau_2 : U[40, 150]$ | 105 |
| Fig. 5H (Left) | $\tau : U[0, 150]$ | 105 |
| Fig. 6B | $\tau_1 : U[0, 60], \tau_2 : U[40, 150]$ | 150 |
| Fig. 6A | $\tau : U[0, 150]$ | 150 |

On subsequent iterations, the same steps are repeated but with parameters drawn from a proposal distribution and with an updated error threshold. On each iteration, the error threshold is set at the first quartile of the accepted sample distances from the previous iteration. The proposal distribution is computed for each iteration *ξ* as a mixture of Gaussian distributions based on the prior distribution and the accepted samples *θ_r_, r* = 1,…, *N* from the previous iteration:

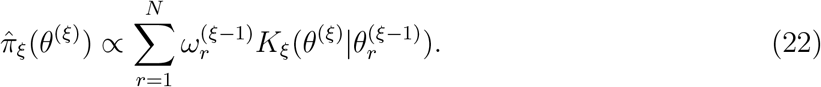

Here 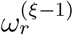 is the importance weight of the accepted sample 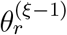 from the previous iteration

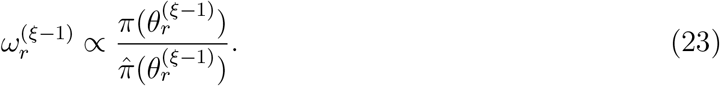

*K_ξ_* is the random walk kernel for the population Monte Carlo algorithm, which is a multivariate Gaussian with the mean 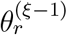 and the covariance equal to twice the covariance of all accepted samples from the previous iteration Σ = 2Cov[*θ*^(*ξ*–1)^]:

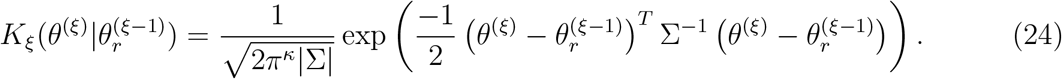

Here *κ* is the number of fitted parameters, and |Σ| is the determinant of Σ.

The convergence of the algorithm is defined based on the acceptance rate accR, which is the number of accepted samples divided by the total number of drawn samples on each iteration. The algorithm terminates when the acceptance rate reaches accRmin, which is set to *accR*_min_ = 0.003 in our simulations. Smaller *accR*_min_ leads to a smaller final error threshold (Fig. 3, right) and a better approximation of the posterior distributions, but requires longer fitting time. To find the MAP estimates, we smooth the final joint posterior distribution with a multivariate Gaussian kernel and find its maximum with a grid search.

### 4.3 abcTau: A Python package for timescales estimation and model comparison with aABC

We developed the abcTau Python package implementing our aABC framework for estimation of timescales from autocorrelations or PSDs of various types of experimental data, and the Bayesian model comparison between different hypotheses. We also provided tutorials as Jupyter Notebooks and example Python scripts to make our framework easily accessible for researchers in different fields.

The minimal requirements for using this package are Python 3.7.1, Numpy 1.15.4 and Scipy 1.1.0. For visualization, Matplotlib >= 3.0.2 and Seaborn >= 0.9.0 are required. The basis of aABC algorithm in the package is adopted from a previous implementation originally developed in Python 2 [82]. Since all parameters of our generative models are positive and sometimes subject to additional conditions (e.g., *τ*_2_ > *τ*_1_), we introduced constraints on sampling from proposal distributions. Moreover, we enhanced the algorithm for parallel processing required for analyzing large datasets.

The abcTau package includes various types of generative models that are relevant for different types data and various methods for computing the autocorrelation or PSD. Using this functionality, users can apply our framework to their time-series data, supplied in a Numpy array structured as trials times time-points. The object oriented implementation of the package allows users to easily replace any function, including generative models, summary statistic computations, distance functions, etc., with their customized functions to better describe statistics of the data. Users can also add their customized generative models directly to the package to create a larger database of generative models available for different applications.

The package also includes a module for Bayesian model selection. This module computes the cumulative distribution of distances from estimated posterior distributions (i.e., Bayes factor for different error thresholds), runs the statistical tests and suggests the best hypothesis describing the underlying processes in data.

#### 4.3.1 Pre-processing for evaluation of direct fit quality

Since Bayesian inference of a full posterior distribution can be computationally expensive, we implemented a fast pre-processing function that uses parametric bootstrapping to determine whether the direct exponential fit provides satisfactory estimates of timescales (Supplementary Fig. 14). In this function, a generative model (e.g., based on a mixture of OU processes) with parameters obtained from the direct exponential fit can be used to generate multiple synthetic datasets, each with the same amount of data as in the original data. For each synthetic dataset, timescales are estimated by direct exponential fitting. The obtained distribution of timescales from this bootstrapping procedure can be compared to the initial direct-fit estimate from the original data. The error between the mean of bootstrapping distribution and the initial direct fit is used to approximately evaluate the direct fit quality. If the error is small enough, the direct exponential fit may be sufficiently accurate. The accuracy of timescale estimates with the direct fit can be further improved by empirical bias-correction using the measured deviation between the mean of the parametric bootstrap distribution and the direct-fit estimates (Supplementary Fig. 5). However, this method does not guarantee an accurate bias correction since the deviation of the direct fit from the ground truth can be larger than the observed deviation between the bootstrap and direct fit. Hence, we recommend users to be conservative with the decision to rely on the direct-fit estimates if accurate estimates of timescales are desired.

### 4.4 Description of neural recordings

Experimental procedures and data pre-processing were described previously [77]. In brief, a monkey was trained to fixate a central dot on a blank screen for 3 s on each trial. Spiking activity was recorded with a 16-channel micro-electrode array inserted perpendicularly to the cortical surface to record from all layers in the visual area V4. For fitting, we used a recording session with 81 trials. We pooled the activity across all channels and calculated the population spike-counts in bins of 1 ms. First, we subtracted the trail-averaged activity (PSTH) from spike-counts to remove the slow trends locked to the trial onset [3]. Then, we computed the autocorrelation of spike-counts using Eq. 2 for each trial and averaged over trials’ autocorrelations.

### 4.5 Parameters of simulations and aABC fits in figures

For all fits, the initial error threshold was set to *ε* = 1. The aABC iterations continued until accR ⩽ 0.003 was reached. All datasets (except for the branching network) consisted of 500 trials each of 1 s duration. The dataset for the branching network (Fig. 4) consisted of 100 trials with 500 time-steps. The parameters for simulations and aABC fits are given in Table 1 and Table 2, respectively.

## Data and Code availability

The abcTau Python package with example data and tutorials are available on GitHub at: https://github.com/roxana-zeraati/abcTau.

## Acknowledgments

This work was supported by a Sofja Kovalevskaja Award from the Alexander von Humboldt Foundation, endowed by the Federal Ministry of Education and Research (RZ, AL), **SMART***START2* program provided by Bernstein Center for Computational Neuroscience and Volkswagen Foundation (RZ), NIH grant R01 EB026949 (TAE), and the Pershing Square Foundation (TAE). We acknowledge the support from the BMBF through the Tübingen AI Center (FKZ: 01IS18039B) and International Max Planck Research School for the Mechanisms of Mental Function and Dysfunction (IMPRS-MMFD). We thank N. A. Steinmetz and T. Moore for sharing the electrophysiological data, which are presented in Ref. [77] and are archived at the Stanford Neuroscience Institute server at Stanford University.

## Author contributions

RZ, TAE and AL designed the research, discussed the results and wrote the paper. RZ wrote the computer code, performed simulations, and analyzed the data.

## 5 Supplementary tables

**Supplementary Table 1.** Running time of the aABC algorithm using the PSD as the summary statistic and 20 parallel processors (Intel(R) Xeon(R) Gold 6152 CPU @ 2.10GHz). Trial duration is 1 sec.

| Generative model | Number of trials | Number of posterior samples | Run time (min) |
| --- | --- | --- | --- |
| One-timescale OU (1 parameter) | 100 | 100 | 5 |
| One-timescale OU (1 parameter) | 500 | 100 | 20 |
| One-timescale OU (1 parameter) | 100 | 500 | 23 |
| Two-timescale OU (3 parameters) | 100 | 100 | 6 |
| One-timescale OU and two oscillations (5 parameters) | 100 | 100 | 7 |

## 6 Supplementary figures

**Supplementary Fig. 1.**
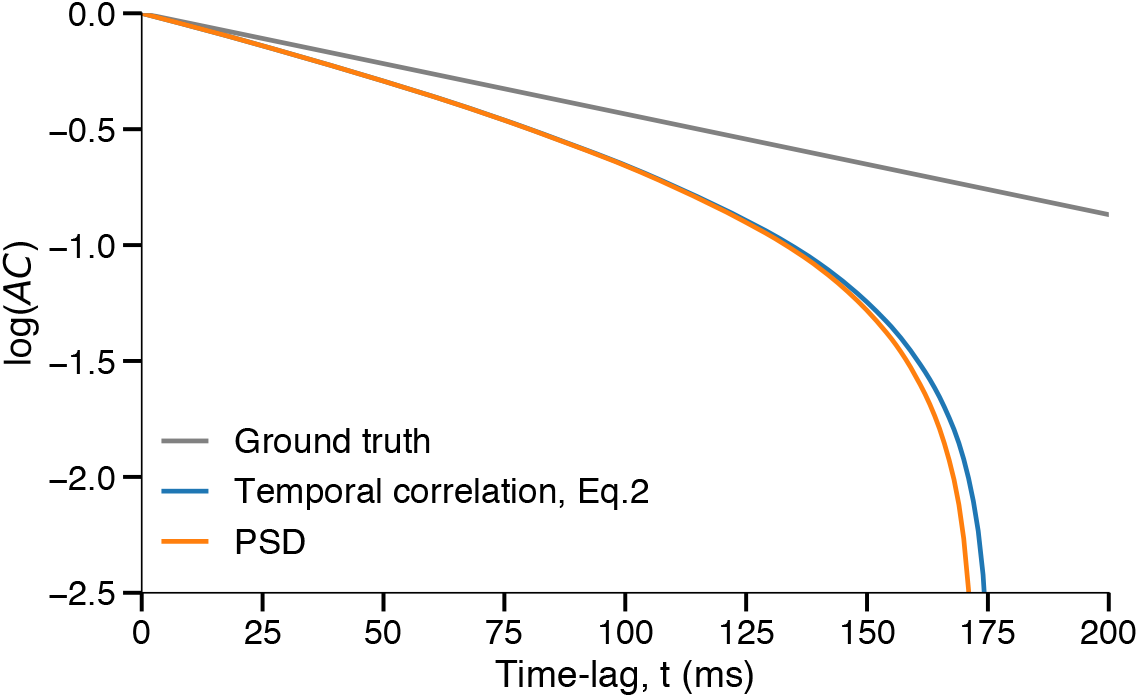
Different methods of estimating the sample autocorrelation from finite data produce biased results. Computing autocorrelations in the time domain (Eq. 2, blue) or via the inverse Fourier transform of the PSD (orange) both produce biased results. Data are generated from an OU process with the timescale *τ* = 100 ms and duration *T* = 1 s. Autocorrelations are averaged over 500 trials (realizations). PSDs are computed after applying a Hamming window to each trial.

**Supplementary Fig. 2.**
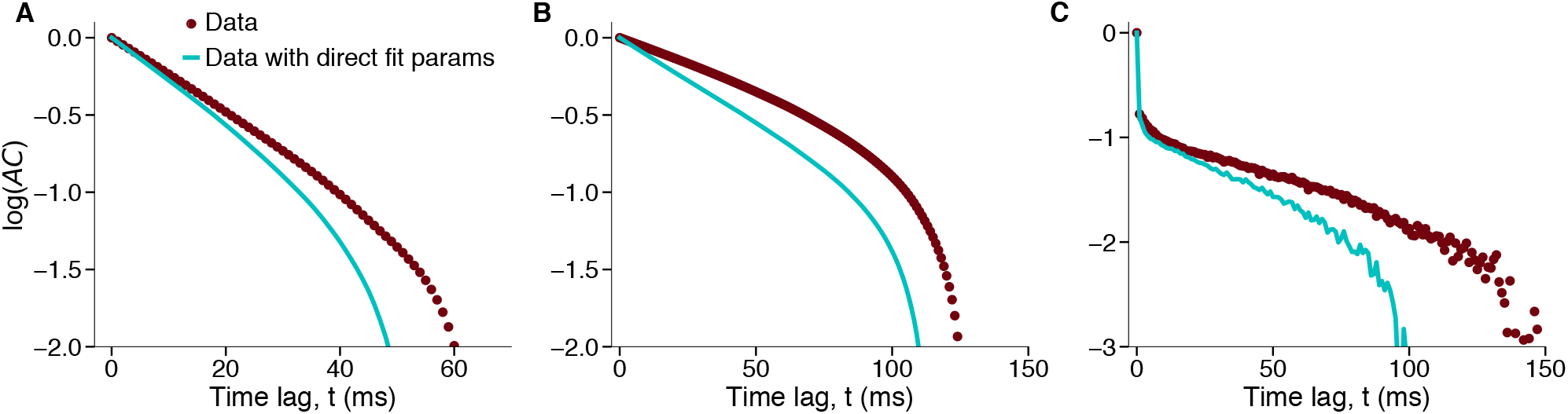
Autocorrelation of synthetic data (cyan) generated with parameters estimated by direct fit deviates from the autocorrelation of the original fitted data (brown). The same synthetic data as in Fig. 1, middle, from A-C respectively. The synthetic data are generated from the correct functional form of generative models with the same statistics (e.g., duration and number of trials) as in the observed data, using parameters obtained from direct exponential fits to observed data autocorrelations.

**Supplementary Fig. 3.**
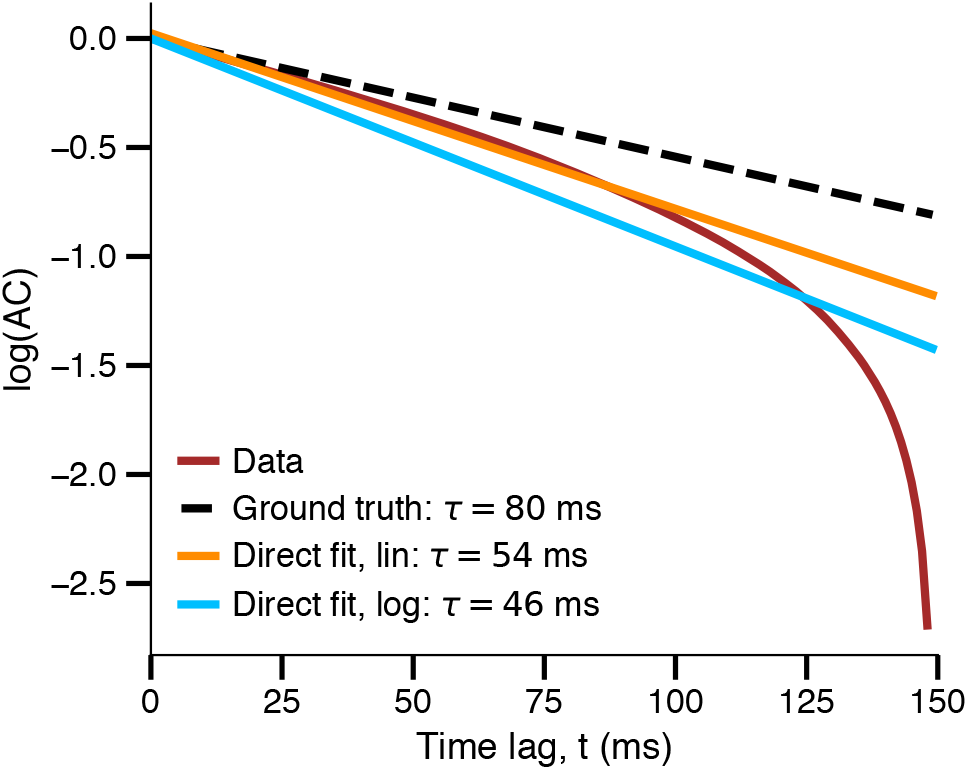
Directly fitting the autocorrelation shape in logarithmic-linear space leads to larger estimation errors than in linear space. The estimated timescale from fitting the sample autocorrelation shape with an exponential decay function is more biased when using logarithmic distances (blue line) than using linear distances (orange line), since it puts more emphasis on the autocorrelation tail which is more affected by the bias (larger deviations between the autocorrelations of data and the ground truth). For a fair comparison between logarithmic and linear fits, only positive correlation values are included in the fitting. Data consists of 500 trials that are generated from an OU process with the timescale *τ* = 80 ms and duration *T* = 1 s.

**Supplementary Fig. 4.**
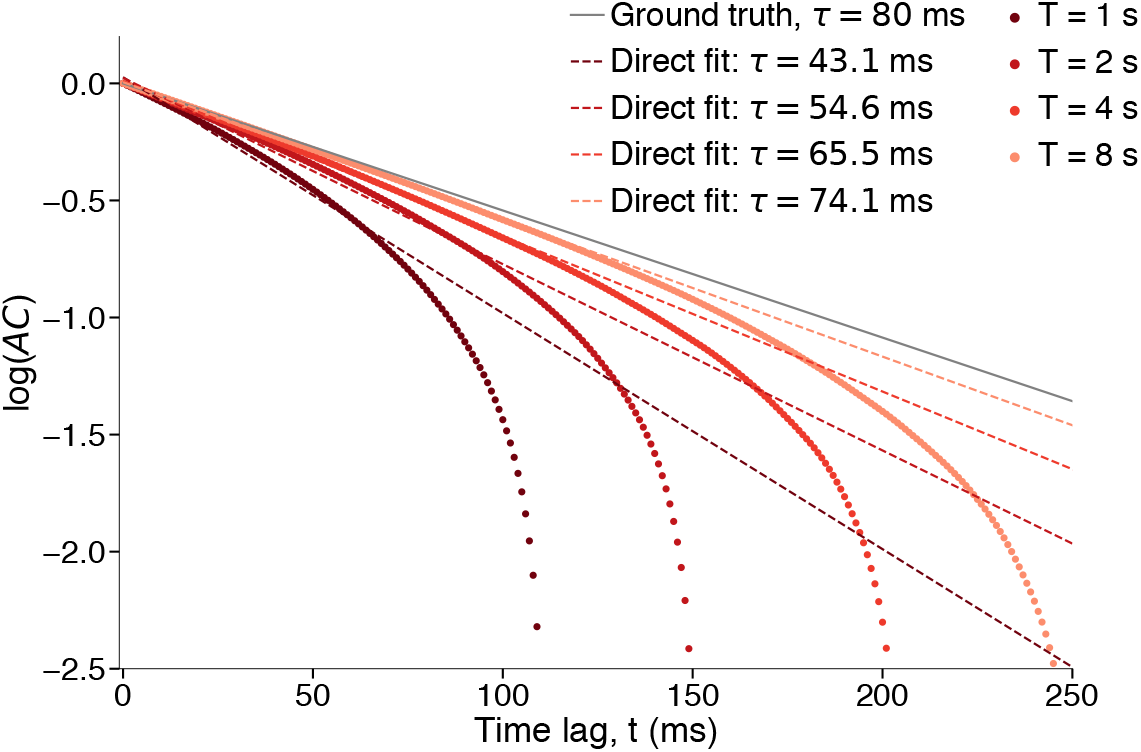
Direct exponential fit produces different estimated timescale for datasets from the same generative process but with different trial duration. Each dataset is generated from an OU process with *τ* = 80 ms and 500 trials each with duration T. Dots - sample autocorrelation, dashed lines - direct exponential fit, solid line - ground truth.

**Supplementary Fig. 5.**
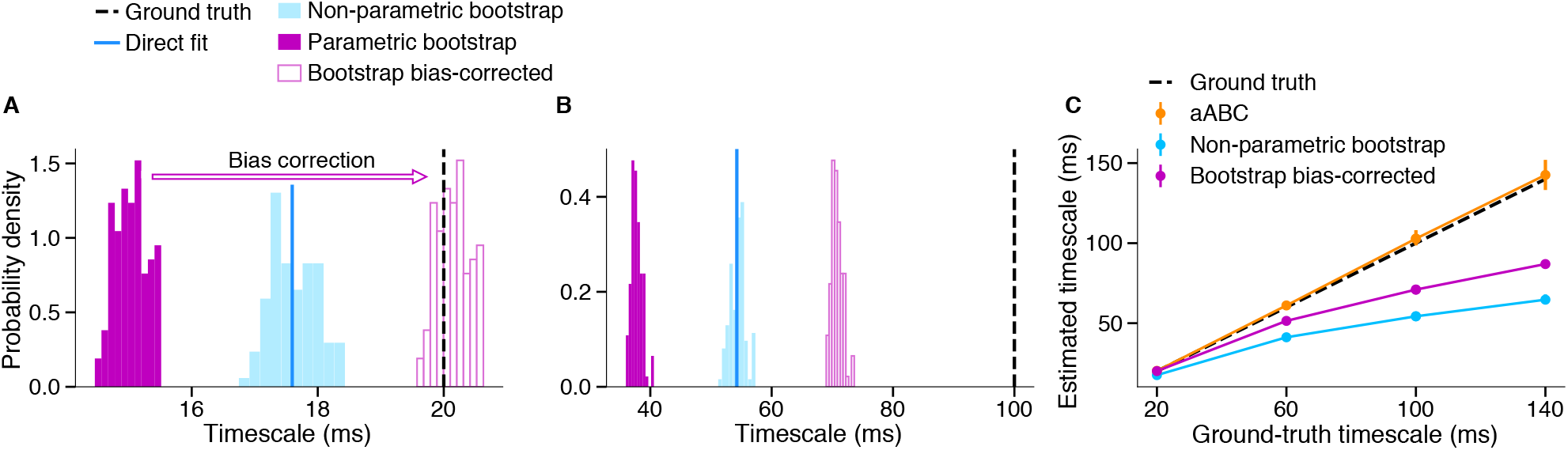
Bootstrapping for the uncertainty estimation and bias correction. **(A-B)** Using bootstrapping for the bias correction does not guarantee unbiased estimation of timescales, since the amount of bias depends on the ground-truth timescale. Data are generated from an OU process with a given ground-truth timescale (dashed line, 500 trials with *T* = 1 s duration). Directly fitting the average autocorrelation shape with an exponential decay function underestimates the timescale (blue line). For parametric bootstrapping, we generate 100 synthetic datasets from an OU process with the timescale estimated by the direct fit and other statistics (e.g., duration and number of trials) as in the original data. We estimate the timescale from each synthetic dataset again with a direct exponential fit (magenta filled bars distribution). The difference between the mean of bootstrapping distribution and the original direct fit estimate can be used as an offset to correct the bias (magenta empty bars distribution). The variance of bootstrapping distribution provides a measure of uncertainty. However, this method only works for some cases (A) and does not guarantee an unbiased estimate for every case (B). For non-parametric bootstrapping, we resample trials from the original data (500 trails with replacement) and estimate the timescale from the average autocorrelation of resampled data with a direct exponential fit (blue distribution). **(C)** Average values of timescales estimated by direct fit with non-parametric bootstrap (blue) and parametric bootstrap with bias correction (magenta) deviate from the ground-truth timescale (dashed lines) and the uncertainty intervals (5 and 95 percentiles, error bars) do not contain these ground-truth values for every example. In contrast, the aABC method provides a reliable mean estimate with uncertainty intervals including the ground-truth timescale (orange). Data are generated from an OU process with the given groundtruth timescales and duration *T* = 1 s (500 trials). Bootstrapping and aABC distributions contain 100 samples. For bootstrapping, the error bars are generally smaller than symbol for the average value.

**Supplementary Fig. 6.**
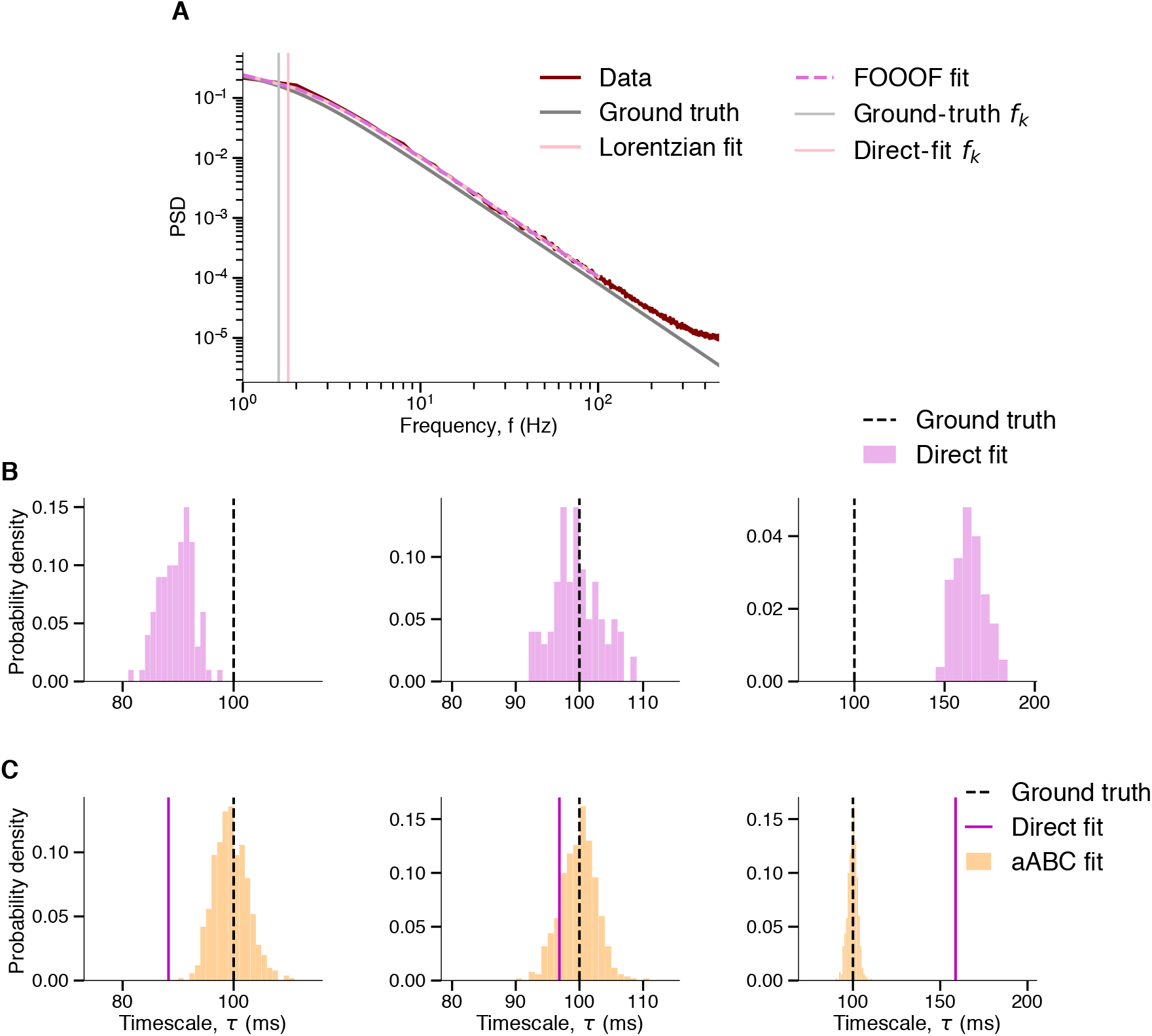
Direct fitting of sample PSD results in biased estimates of timescales, whereas aABC produces accurate estimates. **(A)** The shape of sample PSD (brown) deviates from the ground-truth PSD (gray). Knee frequencies estimated by directly fitting the sample PSD with a Lorentzian function (pink) or using the FOOOF toolbox [42] (magenta) are similar and overestimate the ground-truth knee frequency (i.e. underestimate the ground-truth timescale). FOOOF estimates the timescale by fitting the shape of log(PSD(f)) with the function *b* – log(*k* + *f^χ^*), where *f* is the frequency and *b, k* and *χ* are free parameters. For an exponential decay function *χ* = 2, but FOOOF allows for *χ* ≠ 2. The sample PSD is fitted between 1 Hz and 100 Hz frequencies. **(B)** Timescales estimated in the frequency domain strongly depend on the fitted frequency range and can overestimate, underestimate or produce accurate results depending on the selected frequency range. Distribution of point estimates for the timescales estimated with the FOOOF fit of PSD for 100 independent datasets of the OU process. For each dataset, we averaged the PSD over 500 trials with 1 s duration and the sampling frequency of 1000 Hz. The PSDs are computed using a Hamming window applied to each trial. FOOOF is fitted on the average PSD for each dataset over the frequency ranges of [1, 100] Hz, [1, 200] Hz and [1, 400] Hz from left to right. **(C)** The aABC method estimates the correct timescales (orange posterior distributions) independent of the selected frequency range. Data are the same as in A and the frequency ranges for fitting are the same as B. The direct-fit estimates (magenta vertical lines) are computed using FOOOF.

**Supplementary Fig. 7.**
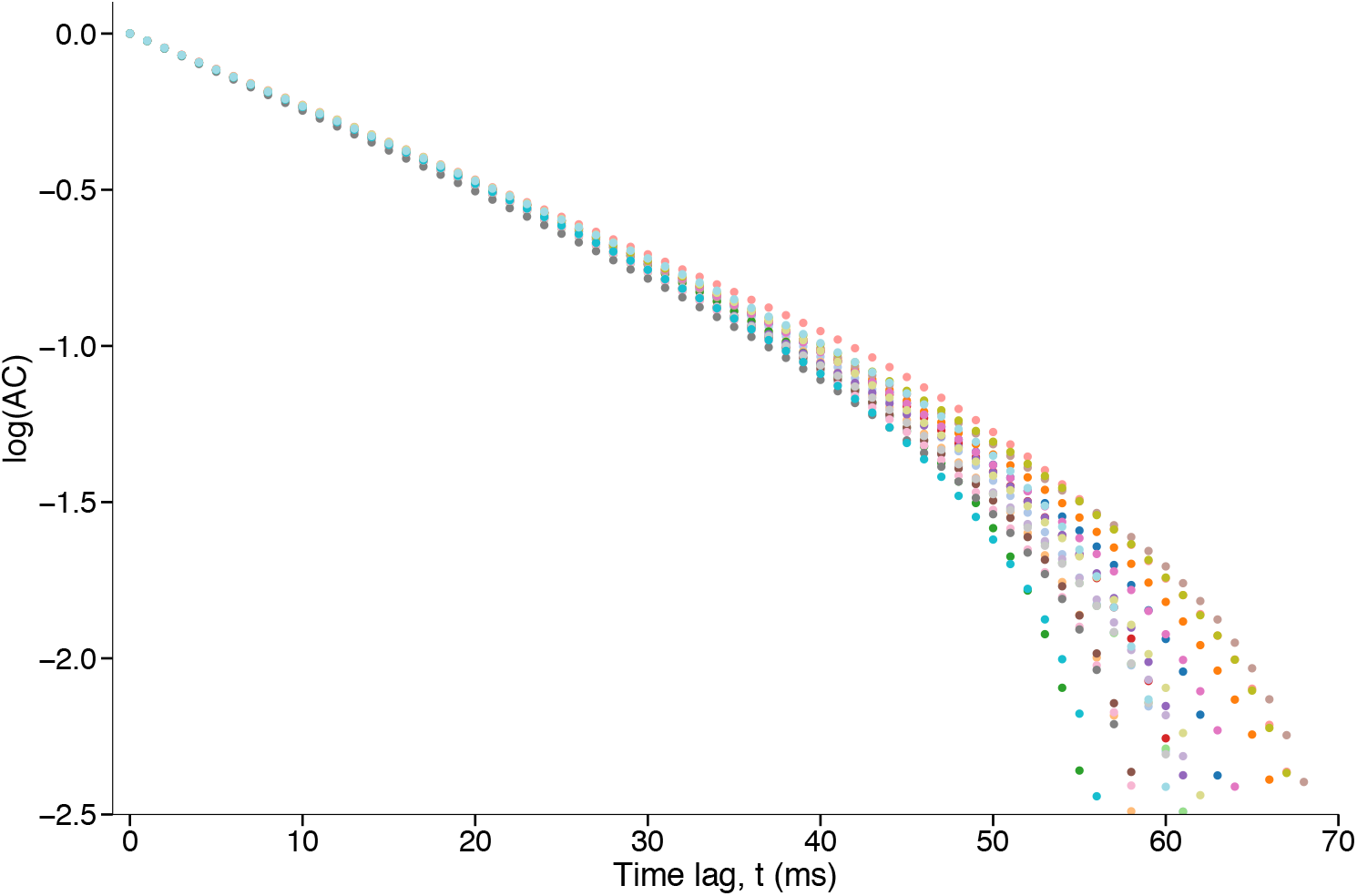
The shape of sample autocorrelation is slightly different across realizations of the same generative process due to noise in the data. Each trace is the average autocorrelation of 500 trials with *T* = 1 s generated from an OU process with *τ* = 20 ms.

**Supplementary Fig. 8.**
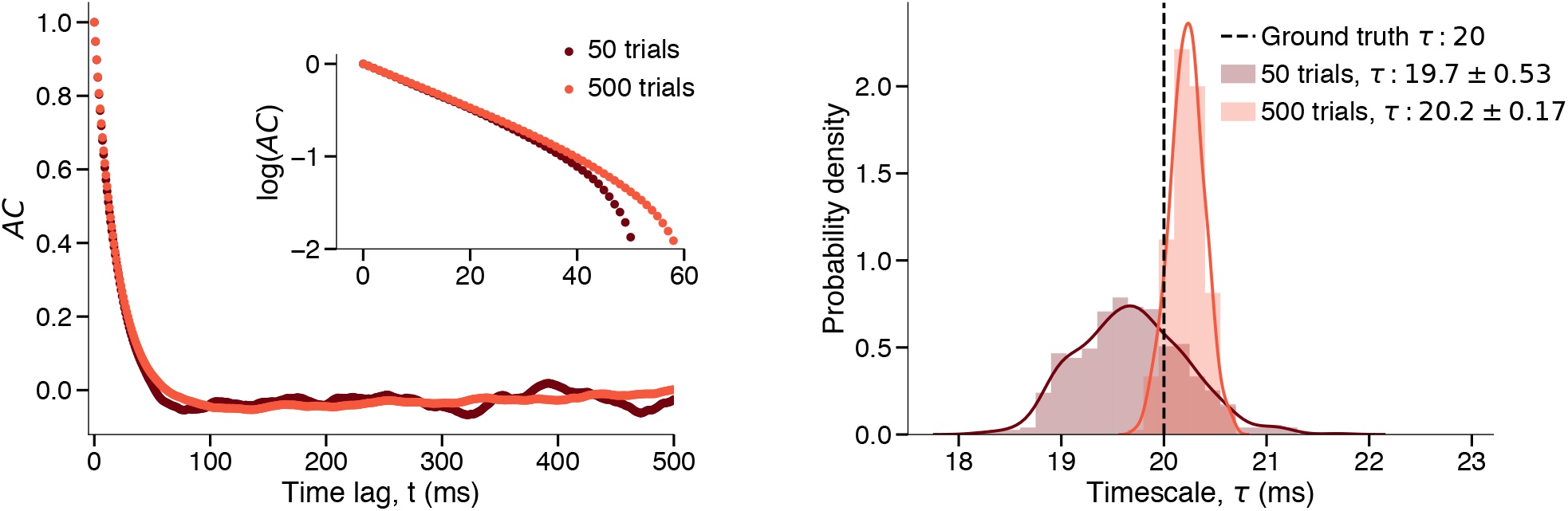
Variance of posterior distributions estimated with aABC captures the estimation uncertainty and depends on the signal to noise ratio. Increasing the number of trials in observed data reduces the noise in sample autocorrelation **(left)** and the estimation uncertainty (narrower posterior distribution) **(right)**. Estimated *τ*: mean ± std. The data was generated from an OU process with *τ* = 20 ms and consisted of trials with *T* = 1 s.

**Supplementary Fig. 9.**
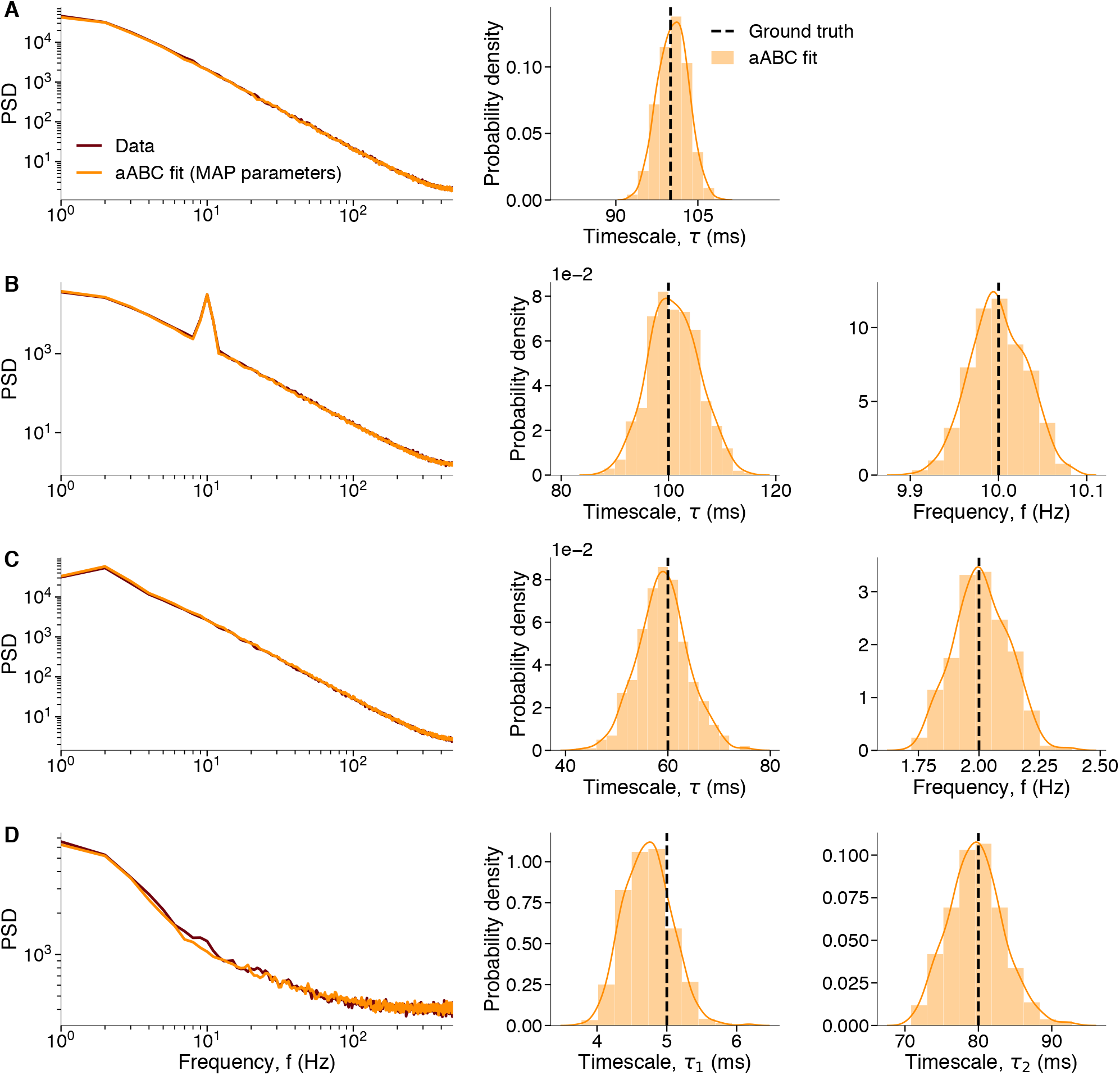
Using PSD in the aABC algorithm accurately recovers the groundtruth parameters. **(A)** Data are generated from an OU process with the timescale *τ* = 100 ms. **Left:** The shape of the data PSD (brown) is accurately reproduced by the PSD of synthetic data from the generative model with the MAP estimate parameters (orange). **Right:** The marginal posterior distributions (histograms overlaid with Gaussian kernel smoothing) recover the ground-truth parameters. **(B)** Same as A for data from a linear mixture of an OU process with the timescale *τ* = 100 ms and an oscillation with frequency *f* = 10 Hz. **(C)** Same as B with *τ* = 100 ms and *f* = 2 Hz. The slow oscillation does not manifest in any clear peak in PSD due to short duration of trials (T = 1 s) and the low frequency resolution. However, aABC algorithm can still uncover the oscillatory component with the correct parameters. **(D)** Same as A for data from an inhomogeneous Poisson process with two timescales *τ*_1_ = 5 ms and *τ*_2_ = 80 ms.

**Supplementary Fig. 10.**
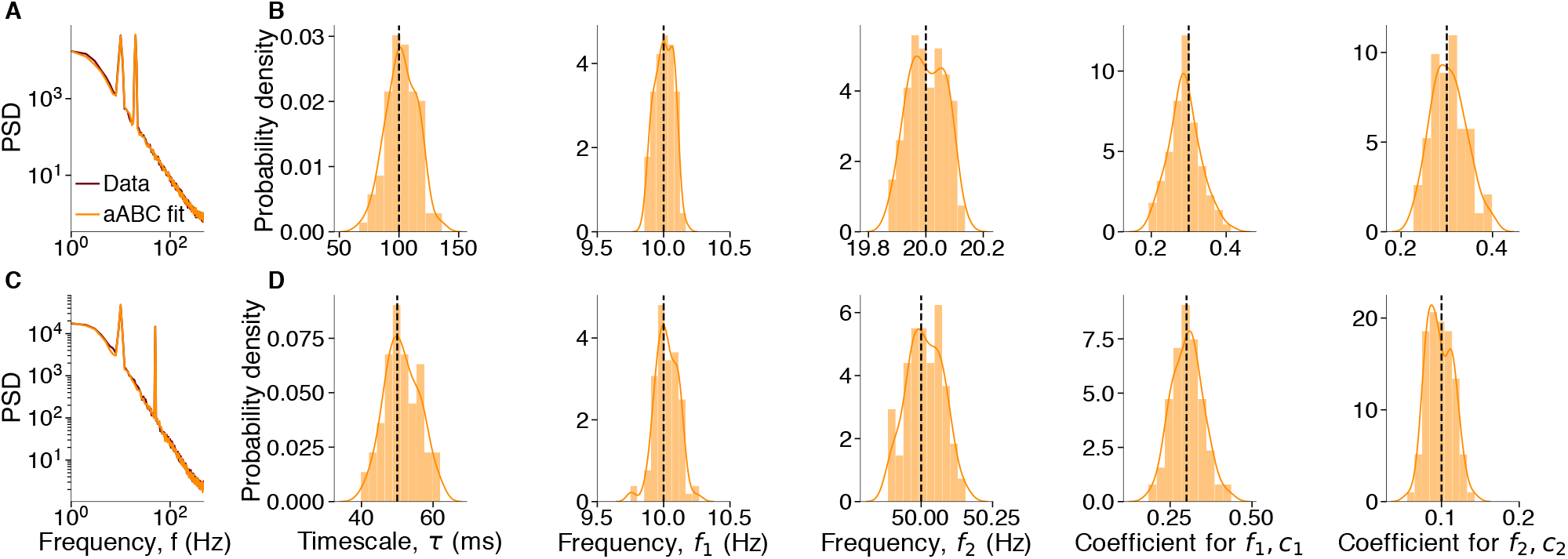
The aABC algorithm can accurately recover the ground-truth parameters even in presence of multiple oscillatory components. **(A-B)** Data are generated from a linear mixture of an OU process with the timescale *τ* = 100 ms and two oscillations with frequencies *f*_1_ = 10 Hz and *f*_2_ = 20 Hz with coefficients *c*_1_ = *c*_2_ = 0.3. **(A)** The shape of the data PSD (brown) is accurately reproduced by the PSD of synthetic data from the generative model with the MAP estimate parameters (orange). **(B)** The marginal posterior distributions (histograms overlaid with Gaussian kernel smoothing) recover the ground-truth parameters (vertical dashed lines). **(C-D)** Same as A-B with *τ* = 50 ms, *f*_1_ = 10 Hz, *f*_2_ = 50 Hz, *c*_1_ = 0.3 and *c*_2_ = 0.1.

**Supplementary Fig. 11.**
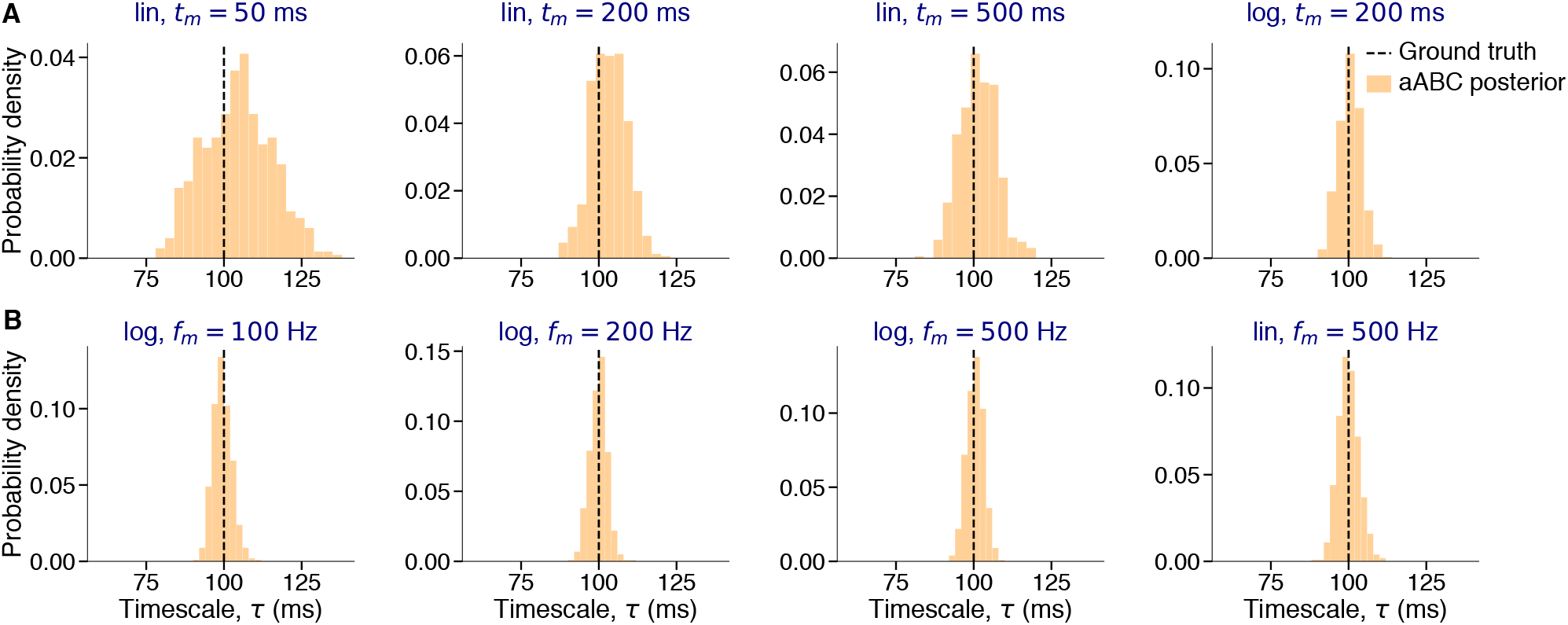
Posterior distributions approximated by the aABC method recover the ground-truth timescales independent of the selected summary statistic and fitting range. Data are generated from an OU process with the timescale *τ* = 100 ms. **(A)** The aABC algorithm fitted the data autocorrelation up to the time-lag *t_m_* (indicated in the figure title) using distances computed in linear (lin) or logarithmic (log) scale. **(B)** The aABC algorithm fitted the data PSD from *f* = 1 Hz up to the frequency *f_m_* (indicated in the figure title) using distances computed in linear (lin) or logarithmic (log) scale.

**Supplementary Fig. 12.**
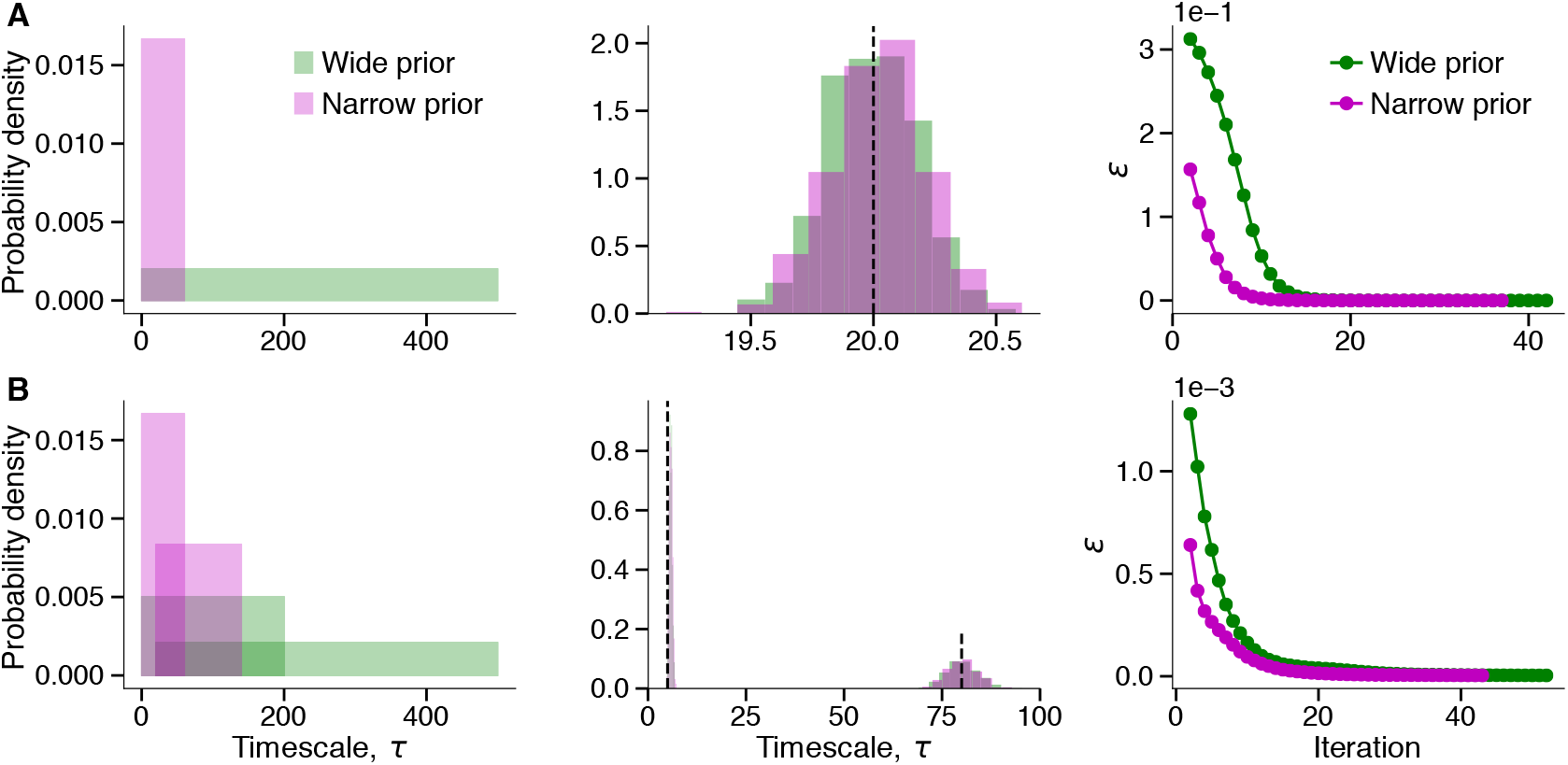
Wide prior distributions for generative model parameters do not affect the shape of posterior distributions and only slow down the fitting procedure. **(A)** Wide and narrow prior distributions (left) give rise to similar posterior distributions (middle), but aABC fit with a narrower prior converges in fewer iterations (right). Data are generated from an OU process with the timescale *τ* = 20 ms and fitted with aABC algorithm and PSD as a summary statistic using two different prior distributions: narrow prior *τ*: *U*[0, 60] and wide prior *τ*: *U*[0, 500] ms, where *U*[·, ·] denotes a uniform distribution. The vertical dashed line indicates the ground-truth timescale. Error thresholds (*ε*) are plotted starting from the second iteration. Initial error threshold for all fits was set to 1. **(B)** Same as A for data generated from an inhomogeneous Poisson process with two timescales *τ*_1_ = 5 ms and *τ*_2_ = 80 ms and prior distributions: narrow prior *τ*_1_: *U*[0, 60] ms, *τ*_2_: *U*[20, 140] ms; wide prior *τ*_1_: *U*[0, 200] ms, *τ*_2_: *U*[20, 500] ms.

**Supplementary Fig. 13.**
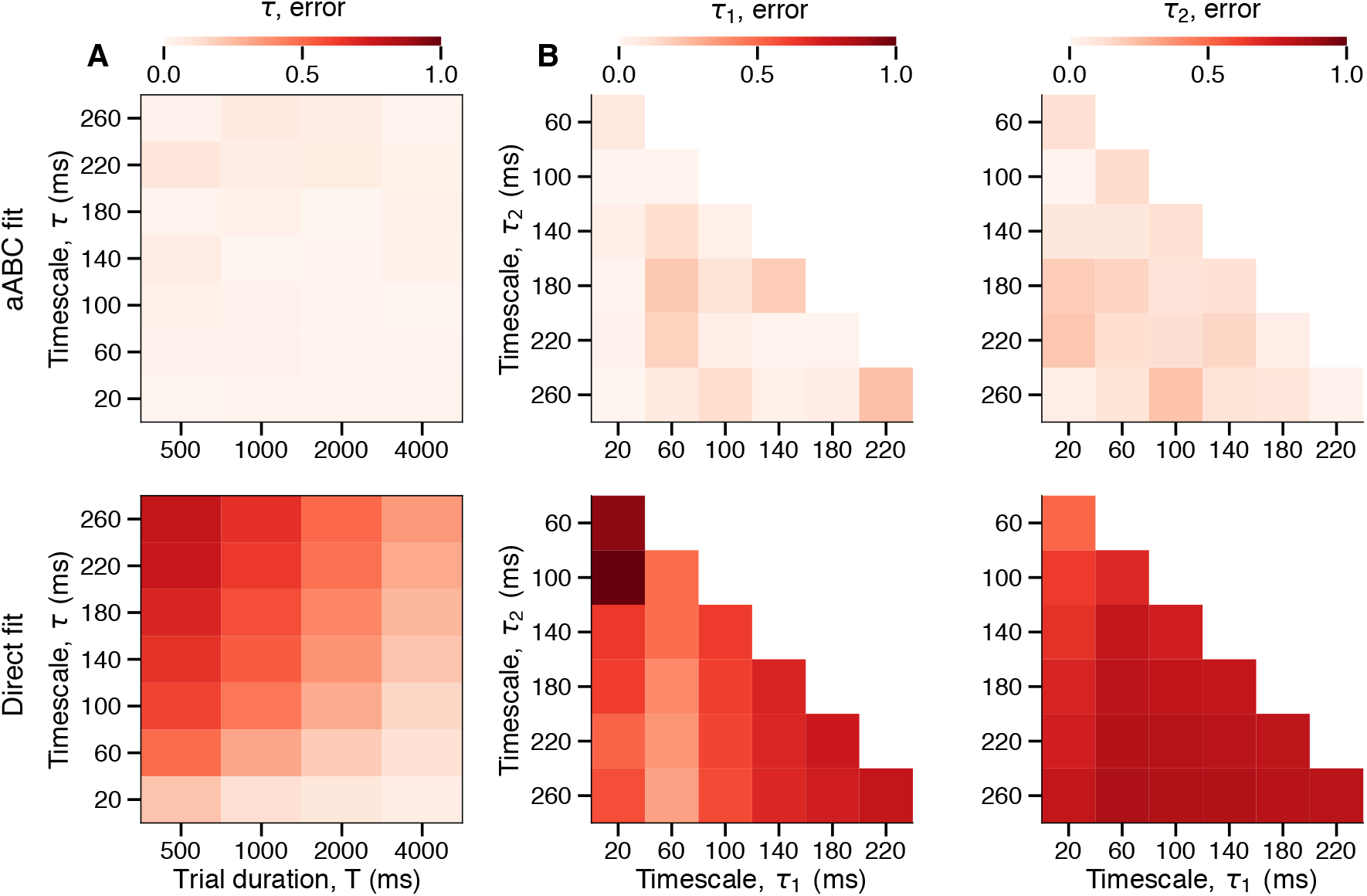
Comparison of goodness of fit for the aABC and direct fitting. **(A)** The estimation error of the MAP estimates from the aABC fit is always small (upper panel), whereas the estimation error of the direct fit depends on the trial duration relative to the ground-truth timescale (lower panel). Data for each fit are generated from an OU process with the given timescales *τ* and the trial duration *T* (500 trials). Estimates of direct fits are computed from fitting the autocorrelation shape with an exponential decay function. Errors are computed as |ground-truth timescale—estimated timescale|/(ground-truth timescale). **(B)** Same as A, for data generated from two-timescale OU processes with the given *τ*_1_ and *τ*_2_ timescales, coefficients *c*_1_ = *c*_2_ = 0.5 and trial duration of *T* = 1000 ms. MAP estimates for both timescales from the joint posterior distribution are always more accurate than direct fitting with the double-exponential function. Here, the relationship between the values of two timescales and accuracy is non-monotonous. In addition, the aABC method returns a full posterior for each estimate (not only the MAP estimate) and the ground-truth timescales are within the bounds of posterior.

**Supplementary Fig. 14.**
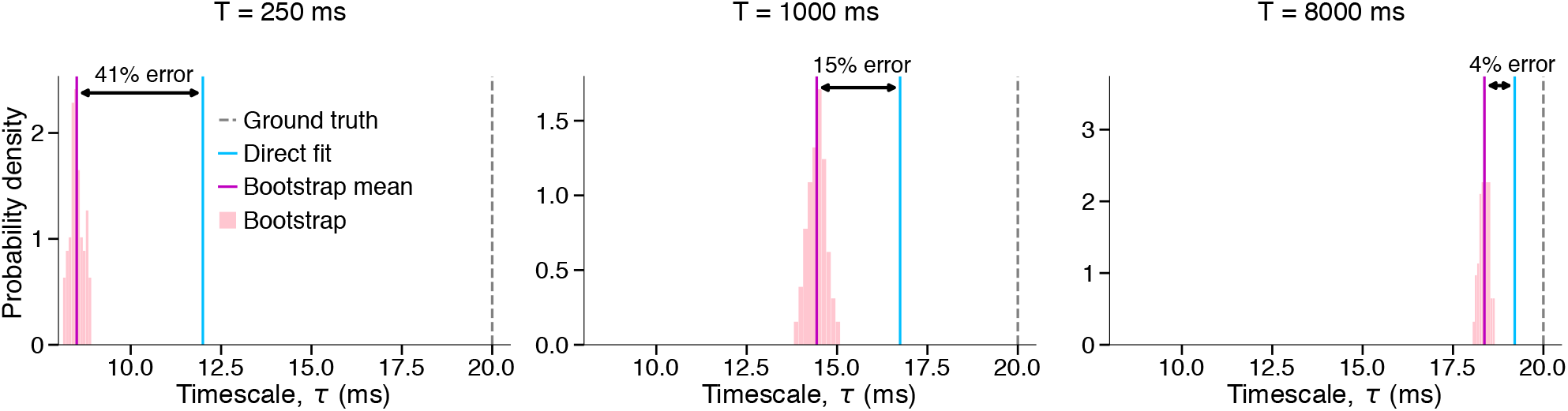
Evaluating the quality of direct fit using abcTau package preprocessing functionality. We use the parametric bootstrapping method to evaluate the quality of direct fit. We generate multiple synthetic datasets from a generative model with parameters estimated by the direct fit and estimate the timescale from each of these datasets again by direct fitting. If the error between the mean of bootstrapping estimates and original direct fit is small enough (based on a user-defined tolerance) the direct fit estimate can be used. Since the bootstrapping error can be smaller than the true error (error between the ground-truth timescale and the direct fit estimate), users should consider setting a stricter tolerance than required. Data in each panel are generated from an OU process with timescale *τ* = 20 ms and the indicated trial duration *T* (500 trials). Increasing the trial duration leads to smaller errors for the direct fit estimates. Errors (black arrows) are computed as |direct-fit estimate – mean(bootstrapping distribution)|/(direct-fit estimate).

**Supplementary Fig. 15.**
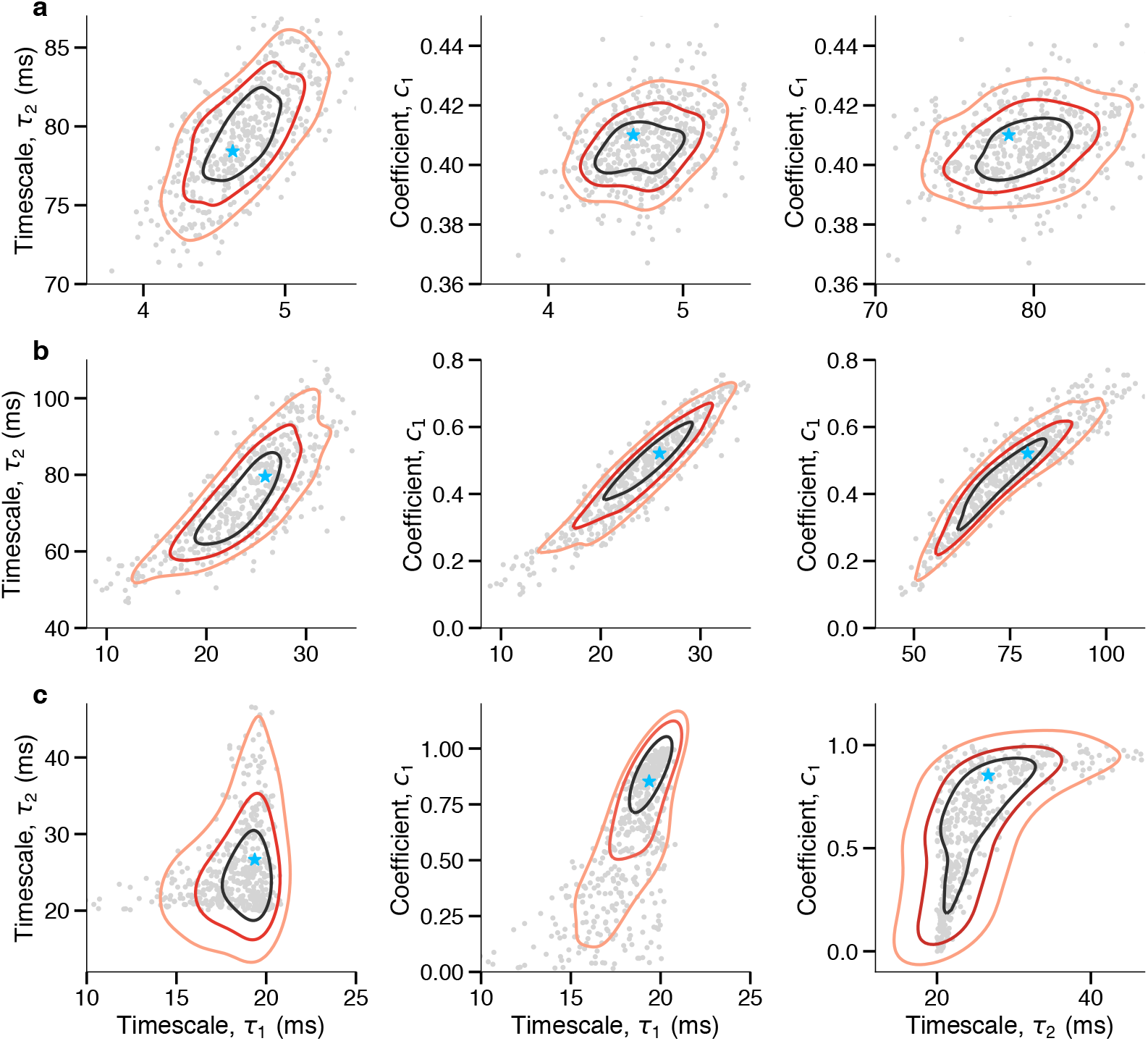
Evaluating the relation between parameters in the joint posterior distributions from the aABC method. **(A-C)** Bivariate marginal distributions obtained from the three-dimensional posterior distribution of parameters from fitting a two-timescale model on different example data (same data as in Fig. 5). Isolines are computed by fitting bivariate Gaussian kernel density on the posterior samples and indicate the parameter combinations with equal probability in the joint posterior (isoline levels are set at *p* = 0.75, 0.5, 0.25 from the inner to outer contours, i.e. for 0.25 isoline 25% of the probability mass will lie within the contour). The blue star shows the MAP estimate from the joint posterior distribution and the gray dots show the samples of the posterior. **(A)** When data has two clear distinct timescales (*τ*_1_ = 5 ms, *τ*_2_ = 80 ms) the joint marginals have a clear peak (black isoline with *p* = 0.75) around the MAP estimate and parameters are more independent from each other (i.e. circular shape of bivariate distributions). **(B)** When the timescales are closer to each other (*τ*_1_ = 20 ms, *τ*_2_ = 80 ms) and the distinction between the timescales is less clear in the autocorrelation shape (Fig. 5C), the parameters become more correlated in the posterior. **(C)** When data has only one timescale (*τ* = 20 ms), there is a degeneracy in parameters of the two-timescale model and fit provides different possible solutions: (i) both timescales are close to the ground truth and the coefficient takes a broad range of values with the same probability, indicating that the coefficient is a sloppy parameter [83], (b) one of the timescales is away from the ground truth and its coefficient is close to zero, suggesting that this additional timescale is a sloppy parameter. This degeneracy indicates that the model is overparametrized. Note that we enforce *τ*_1_ < *τ*_2_ which generates asymmetry in the posterior.

**Supplementary Fig. 16.**
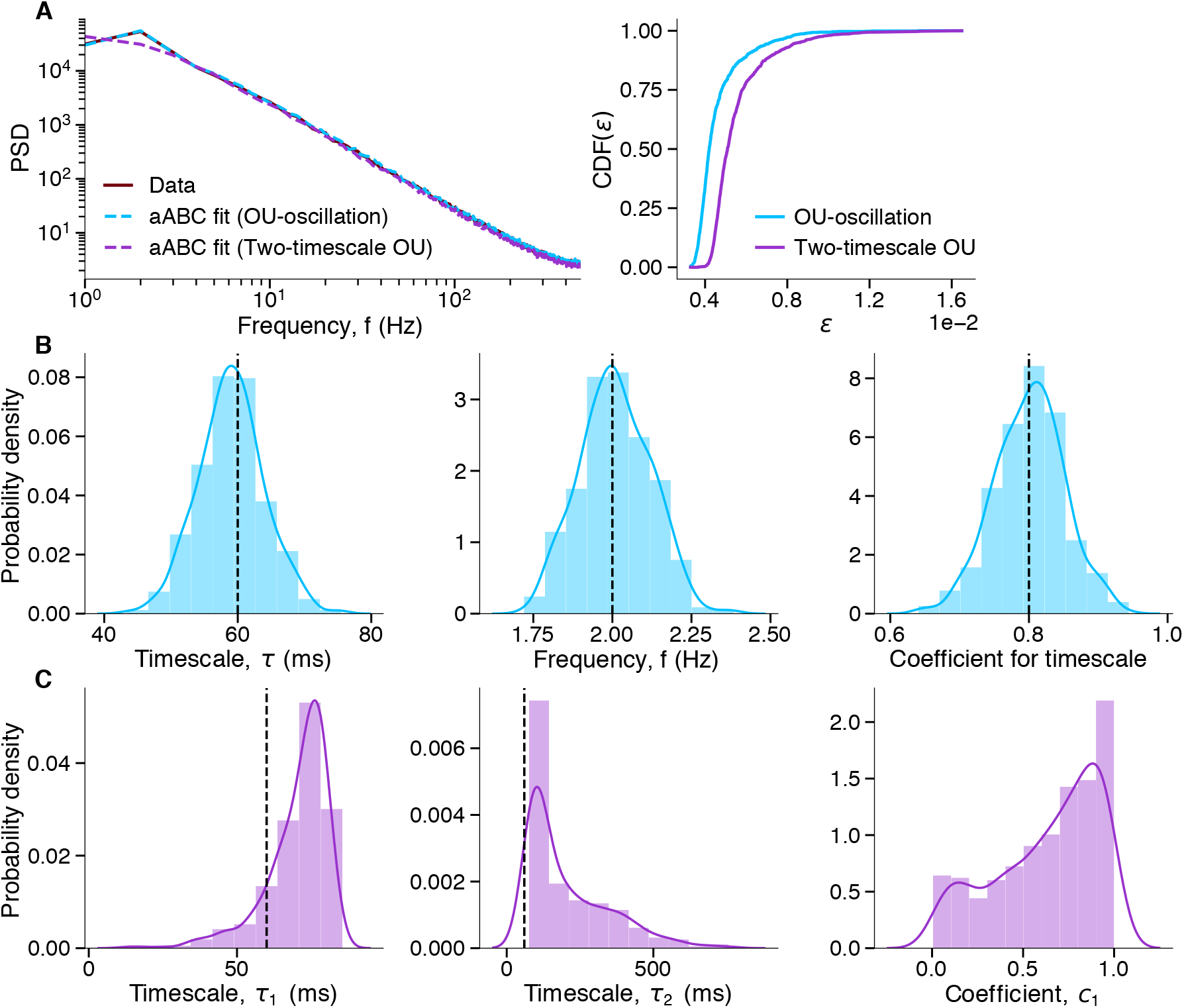
Model selection with ABC discriminates between periodic (oscillatory) and aperiodic (exponential decay) components. **(A-C)** Data are generated from a linear mixture of an OU process with the timescale *τ* = 60 ms and an oscillation with frequency *f* = 2 Hz (same as Supplementary Fig. 9C). Using the ABC model comparison method, we tested whether the shape of the PSD from these data can be equally well described with a two-timescales OU process without any oscillatory components. **(A)** The data PSD is fitted with two models: 1) a linear mixture of an OU process and oscillation (blue), 2) a two-timescale OU process (purple) (left). Since cumulative distribution of distances for the first model is larger than for the second model for all *ε*, the model with the linear mixture of an OU process and oscillation is selected (right). **(B)** The marginal posterior distributions (histograms overlaid with Gaussian kernel smoothing) for the linear mixture of an OU process and oscillation. Vertical lines indicate the ground-truth values. **(C)** The marginal posterior distributions (histograms overlaid with Gaussian kernel smoothing) for the two-timescale OU process. Vertical lines indicate the ground-truth value of the single ground-truth timescale.

**Supplementary Fig. 17.**
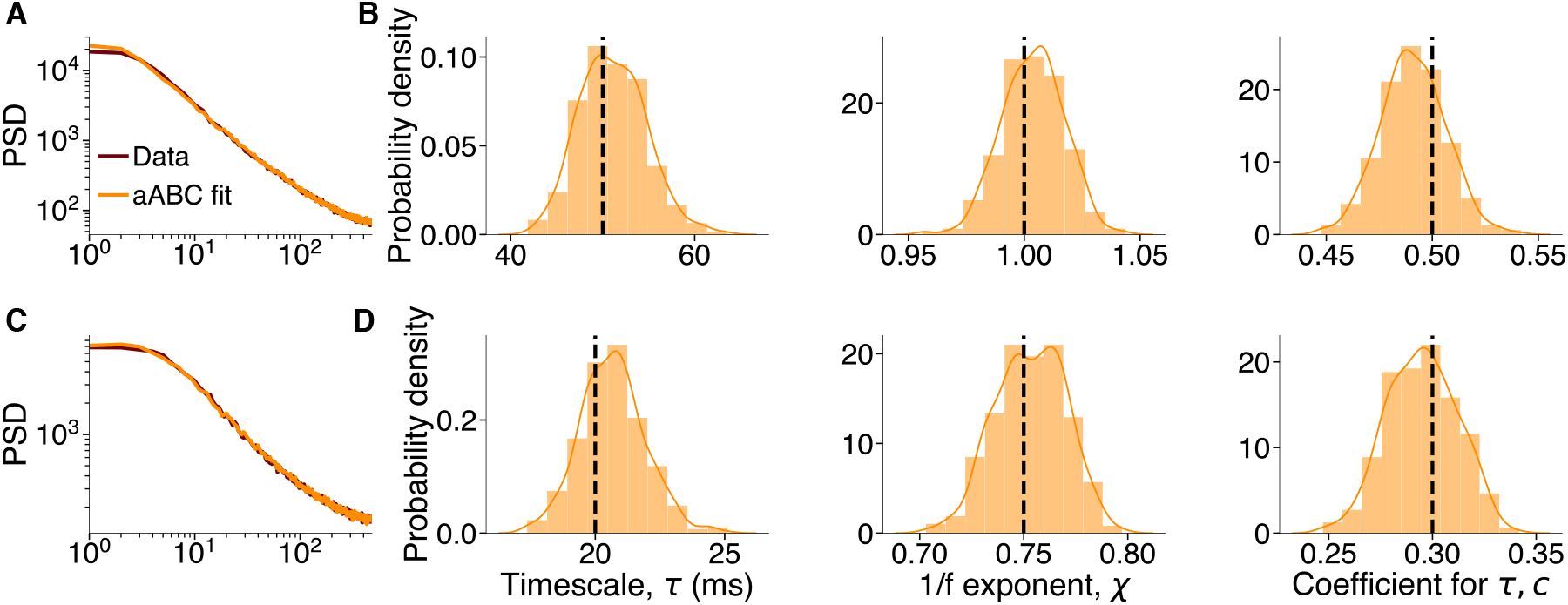
Estimating timescales with aABC in the presence of background processes. **(A-B)** Data are generated from a linear mixture of an OU process with the timescale *τ* = 50 ms (coefficient *c* = 0.5) and random time-series generated from a 1/*f* PSD shape with exponent *χ* = 1 over the frequency range of [1, 500] Hz (corresponding to the time-domain signals with *T* = 1 s duration). **(A)** The shape of the data PSD (brown) is accurately reproduced by the PSD of synthetic data from the fitted generative model with the MAP estimate parameters (orange). The generative model is the same as for the original data (i.e. a linear mixture of an OU process and random time-series generated from a 1/*f* PSD shape). **(B)** The marginal posterior distributions (histograms overlaid with Gaussian kernel smoothing) recover the ground-truth parameters (vertical dashed lines). **(C-D)** Same as C-D with *τ* = 20 ms, *χ* = 0.75, *c* = 0.3.

**Supplementary Fig. 18.**
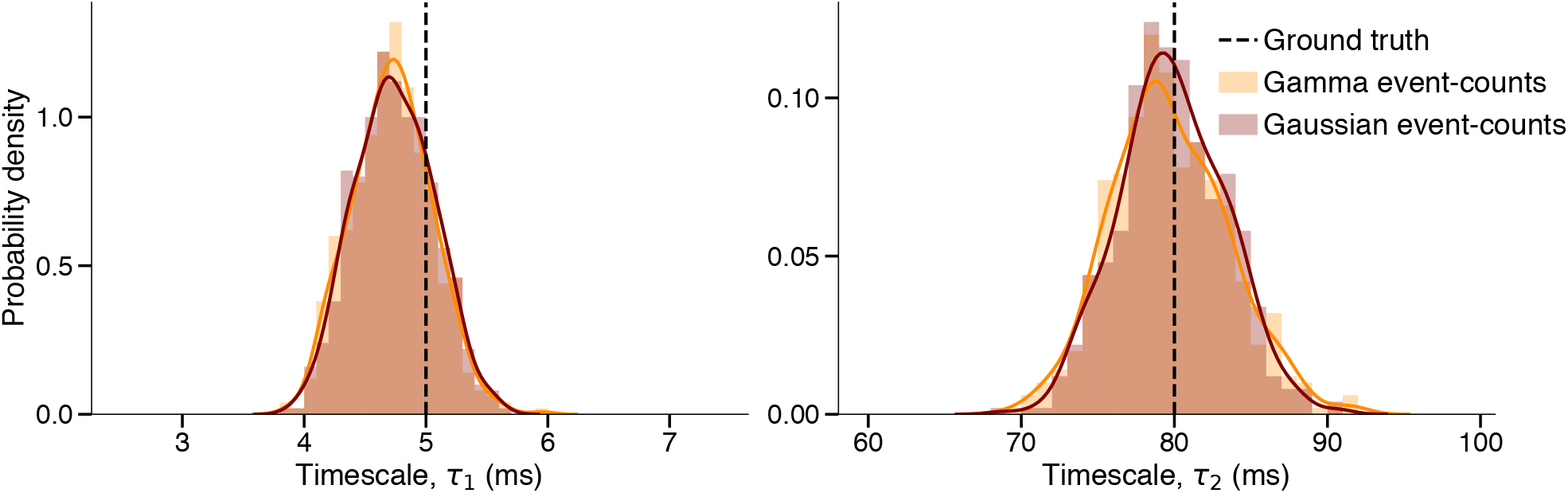
Doubly stochastic generative models with the Gaussian and gamma distributions of event-counts produce similar aABC fits. We fitted the inhomogeneous Poisson process example from Fig. 1C using aABC with generative models which had a Gaussian or gamma event-count distributions with *α* = 1. Both generative models recover the ground-truth timescales and produce posterior distributioins that are not significantly different (Wilcoxon rank-sum test: *P*_*τ*_1__ = 0.23, *P*_*τ*_2__ = 0.37).

**Supplementary Fig. 19.**
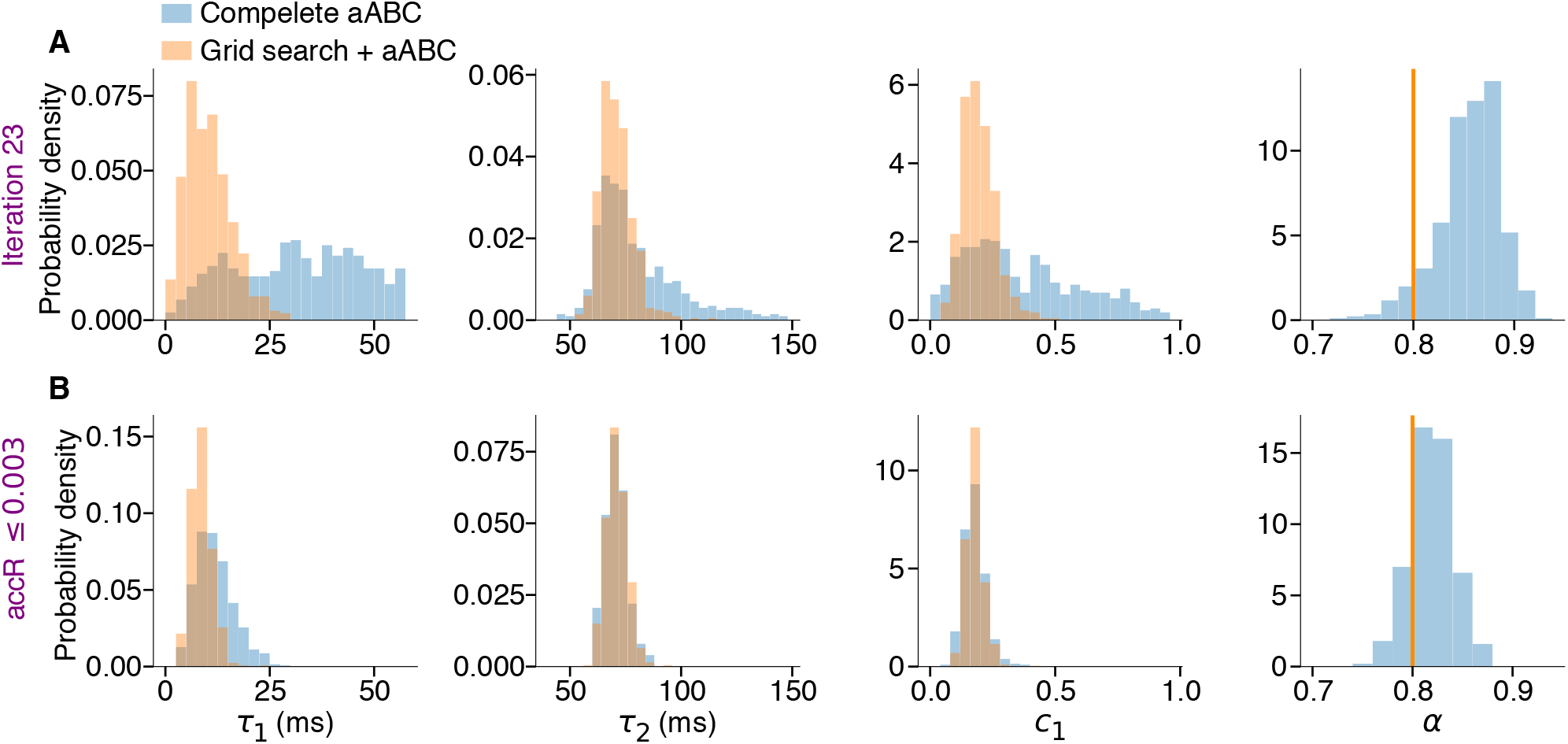
Estimating the parameter *α* of the doubly stochastic generative model with aABC and with the grid search. We fitted the neural data autocorrelation in Fig. 6 considering α as an additional parameter of the generative model estimated with aABC (complete aABC). We compared the results of complete aABC with first estimating α with a grid search and then fitting the rest of parameters with aABC (grid search + aABC). **(A)** Starting from the same error threshold *ε* = 1, after the same number of iterations (23), the complete aABC has a larger variance of marginal posterior distributions than the grid search + aABC. This result indicates that the grid-search method reaches narrower posteriors in fewer iterations. Moreover, for the complete aABC, the posterior of *α* is relatively narrow, while posteriors of all other parameters are wider at iteration 23. This observation shows that the complete aABC fits α first and almost independently from the other parameters of the generative model. **(B)** After running both methods until they reach *accR* ≤ 0.003, MAP estimates with complete aABC converge toward the estimations from the grid search + aABC.

## Notes

https://github.com/roxana-zeraati/abcTau

### Competing Interest Statement

The authors have declared no competing interest.

### Summary of Updates

added more generative model flexibility; extended supplementary materials

